# Alternative catalytic mechanisms driven by structural plasticity is an emerging theme in HAS-GTPases, Era and FeoB

**DOI:** 10.1101/2020.08.16.253419

**Authors:** Sahil Batra, Ashok Kumar, Balaji Prakash

## Abstract

GTP hydrolysis is the underlying basis for functioning of ‘biological switches’ or GTPases. Extensively studied GTPases, Ras and EF-Tu, use a conserved Gln/His that facilitates the activation of attacking water for nucleophilic attack. However, this is insufficient to explain catalysis in <u>H</u>ydrophobic <u>A</u>mino acid <u>S</u>ubstituted (HAS)-GTPases that naturally possess a hydrophobic residue in lieu of Gln/His. We had previously reported a bridging water-chain mediated catalytic mechanism for HAS-GTPase FeoB; which utilizes two distantly-located but conserved glutamates. Curiously, mutating these does not abolish GTP hydrolysis. Similarly, in this study we report our observations on another HAS-GTPase Era, wherein the mutants of catalytically important residues continue to hydrolyze GTP. We attempt to rationalize these inquisitive observations on GTP hydrolysis by FeoB and Era mutants. We propose a general theory that appears common to at least three classes of GTPases, where ‘alternative mechanisms’ emerge when the primary mechanism is disrupted. Based on the analysis of crystal structures of FeoB and Era mutants, bound to the transition state analogue GDP.AlF_x_, this work suggests that in the absence of catalytically important residues, the active site waters in both FeoB and Era undergo re-arrangements, which in turn helps in sustaining GTP hydrolysis. Similar employment of alternative mechanisms was also suggested for the catalytic mutants of hGBP1. Importantly, such alternatives underscore the robustness of GTP hydrolysis mechanisms in these systems, and raise important questions regarding the need for persistent GTP hydrolysis and the physiological relevance of structural plasticity seen here.

## Introduction

G-proteins or Guanosine tri-phosphatases (GTPases) are GTP hydrolyzing enzymes that constitute an important class of regulatory proteins, which moderate crucial cellular processes such as cell growth, signaling, protein synthesis and molecular transport^1–3^. The enzymatic hydrolysis of GTP to GDP (guanosine di-phosphate) and an inorganic phosphate is frequently coupled with remarkable structural rearrangements in ‘switch’ regions (switch-I and switch-II), which form an interface with the downstream effector molecules^3,4^. GTPases attain an active or ‘ON’ state in the presence of GTP, and interact with other effector domains/proteins. GTP hydrolysis toggles it to a GDP bound ‘OFF’ state, curtailing its aforementioned interactions. GTPases effectively communicate a change in the nucleotide state to activate/inactivate biochemical pathways^1,5^, and are hence termed as ‘biological switches’.

GTP hydrolysis is essential for timely inactivation of biological switch. Importantly, mutation(s) that affect GTP hydrolysis, permanently ‘Switch ON’ downstream signaling. The proto-oncogene *ras* codes for a GTPase that regulates cell growth^1^. Mutations that abolish its hydrolytic activity result in a loss of regulation, thus culminating in cancerous growth^6,7^. Genetic analyses identified a conserved glutamine (Gln61) – adjacent to the pivotal G3 motif of switch-II region – to be indispensable for GTP hydrolysis, and found that it is frequently replaced by a hydrophobic residue in many tumors^7,8^. The GTP hydrolysis reaction entails a nucleophilic attack by an activated water molecule on the γ-phosphate, resulting in the cleavage of the phosphoanhydride bond^1,4^. The reaction proceeds through a planar phosphoryl intermediate. Structural biologists have ingeniously exploited the chemical similarity of aluminium fluoride (AlF_x_: AlF_3_/AlF_4_^-^) with phosphoryl intermediate^9,10^ to efficiently crystallize GTPases with GDP.AlF_x_ complex^11–16^. GDP.AlF_x_ complex serves as an excellent transition state analogue of the GTP hydrolysis reaction^17^. The transition state structure reveals that Gln^cat^ juxtaposes the attacking water molecule and γ-phosphate of GTP to accomplish its hydrolysis^13^. Simulations and experimental studies have further established that Ras utilizes ‘<u>S</u>ubstrate <u>A</u>ssisted <u>C</u>atalytic (SAC) mechanism’ for GTP hydrolysis; wherein the terminal phosphate of GTP acts as the general base to activate the nucleophilic water^18–24^. Gln^cat^ facilitates the nucleophile activation by assisting in proton transfer to the terminal phosphate.

Although critical, this Gln^cat^ is itself insufficient to initiate hydrolysis. It is oriented in a catalytically competent conformation through interaction with an externally provided Arg residue – ‘Arginine finger’ – from RasGAP (GTPase activating protein)^13,25^. Furthermore, Arg neutralizes the negatively charged intermediate state, accelerating GTP hydrolysis. Notably, many variations to this catalytic arrangement are possible; with cations assuming the role of Arginine finger^15,26,27^, and His replacing Gln^cat^ – as exemplified by translational GTPase EF-Tu^28–32^. However, despite these variations, these classical GTPases employ SAC mechanism for GTP hydrolysis^23,24,32^. The Gln^cat^ or corresponding residue is consistently supplied from a unique location – adjoining the conserved DxxG G3 motif.

Intriguingly, there exists a class of functional GTPases that harbor a hydrophobic substitution in lieu of Gln^cat^. These <u>H</u>ydrophobic <u>A</u>mino-acid <u>S</u>ubstituted (HAS)-GTPases can efficiently catalyze GTP hydrolysis despite the absence of Gln^cat 33^. Sequence and structural analysis of HAS-GTPases identified frequent insertions in the switch-II loop, and indicated that unlike Gln^cat^ the hydrophobic amino acid would orient away from the active site; thus creating an opportunity for the catalytic residue to be supplied from different directions^33^. The available transition state structures for a few HAS-GTPases (atlastin^34^, MnmE^15^ and dynamin^16^) illustrate distinct spatial origins of the catalytic residue(s) (**Figure S1**). In distinction to Gln^cat^ of Ras, HAS-GTPases utilize a bridging water-mediated interaction with the attacking water (**Figure S1**). HAS-GTPases present divergent catalytic mechanisms and necessitate a thorough investigation to unravel their diversity.

FeoB is a unique multi-domain HAS-GTPase that facilitates Fe^2+^ transport across the inner membrane of bacteria^35,36^. The G-domain present at the N-terminal cytosolic part (NFeoB) regulates Fe^2+^ transport through the C-terminal transmembrane domain^37,38^. Significant insights into the mechanistic basis of cytosolic domain’s function arise from elaborate structural studies performed with *Streptococcus thermophilus* NFeoB (*St*NFeoB)^39–42^. However, catalytically important residue(s) could not be deduced from these structural studies. Strong conservation of the pair - Glu66 and Glu67-across homologues^41^, along with obligate requirement of Glu66 for physiological function^38^, strengthens the possibility of catalytic role for these glutamates. However, biochemical analysis of each of the glutamate mutants, only exhibits lowered GTP hydrolysis, but not a complete abolishment^41^. This raises other possibilities for catalysis in FeoB.

In this study, we attempt to explicitly understand the role of the glutamate pair in GTP hydrolysis. We have determined the transition state structures of *St*NFeoB mutants, and combining these with hybrid quantum mechanical/molecular mechanical (QM/MM) simulations (reported previously^43^) we delineate the effect of each mutation on GTP hydrolysis. Together these studies illustrate the catalytic importance of this glutamate pair and highlight the plasticity exhibited by *St*NFeoB mutants. Interestingly, the unique structural features conserved among HAS-GTPases facilitate this structural plasticity, as observed in *St*NFeoB; this seems to be an outcome of rudimentary catalytic machinery common to all the HAS-GTPases.

Era is a HAS-GTPase that functions in the assembly of 30S subunit^44–46^. The G-domain of Era is followed by a C-terminal K homology (KH) domain, which interacts with 16S rRNA from 30S subunit^44,47,48^. The GTP hydrolysis mechanism of Era, so far, remains elusive. A comparison of the catalytic mechanisms in FeoB and Era would be advantageous to deduce the general principles of GTP hydrolysis in HAS-GTPases. We investigate this aspect by structural and biochemical studies of two Era homologues – *Aquifex aeolicus* Era (*Aa*Era) and *E. coli* Era (*Ec*Era) – to identify catalytically important residues. We again observe that mutation of these residues does not abolish GTPase activity of Era; rather, it activates alternative routes for initiating GTP hydrolysis.

These attempts to unravel the catalytic mechanisms in HAS-GTPases, uncovers their structural plasticity. Importantly, such alternatives to accomplish GTP hydrolysis underscore the robustness of the system.

## Results and Discussions

### Glu66 and Glu67 play catalytic roles for GTP hydrolysis

We wanted to precisely examine the importance of each of these residues E66 and E67 in GTP hydrolysis of *St*NFeoB. Therefore, we attempted to co-crystallize the single mutants E66A and E67A in the presence of GDP.AlF_x_, complexed with K^+^ and Mg^2+^. The crystal of *St*NFeoB E67A diffracted to a resolution of 2.35 Å and belonged to space group P2_1_2_1_2_1_ with two monomers in the asymmetric unit (**Table 1**). The nucleotide binding site of each monomer of this active state structure of E67A mutant contains a GDP.AlF_4_^-^ complex, a magnesium ion (Mg^2+^) and a potassium ion (K^+^) (**Figure S2** and see **Supplementary information**).

**Table 1:** Data collection and refinement statistics.

|  | **StNFeoB E67A** | **StNFeoB E66A.E67A** | **AaEra Y63A** |
| --- | --- | --- | --- |
| **PDB ID** | 7BWV | 7BVU | 7C1O |
| **Data processing** |  |  |  |
| Wavelength (Å) | 1.542 | 1.542 | 1.542 |
| Resolution (Å) | 2.35 | 2.5 | 2.18 |
| Space group | P 2<sub>1</sub> 2<sub>1</sub> 2<sub>1</sub> | P 2<sub>1</sub> 2<sub>1</sub> 2<sub>1</sub> | P 2<sub>1</sub> 2<sub>1</sub> 2<sub>1</sub> |
| Unit cell dimensions |  |  |  |
| *a*, *b*, *c* (Å) | 48.7, 73.61, 156.75 | 48.62, 72.40, 157.96 | 36.38, 72.86, 120.91 |
| $\alpha$, $\beta$, $\gamma$ (°) | 90, 90, 90 | 90, 90, 90 | 90, 90, 90 |
| Total reflections | 327983 | 140774 | 135401 |
| Unique reflections | 24305 (3827) | 18799 (900) | 17467 (2744) |
| Multiplicity | 13.49 (13.21) | 7.5 (5.7) | 7.8 (7.5) |
| Completeness (%) | 99.8 (99.2) | 92.87 (89.48) | 99.8 (99.3) |
| $\langle I/\sigma(I) \rangle$ | 21.51 (3.9) | 20.14 (5.7) | 18.12 (2.14) |
| Wilson B-factor | 35.7 | 36.10 | 42.8 |
| $R_{merge}$ | 0.113 (0.776) | 0.073 (0.273) | 0.077 (0.931) |
| $R_{measure}$ | 0.118 (0.808) | 0.079 (0.301) | 0.082 (0.999) |
| $CC_{1/2}$ (%) | 99.9 (89.0) | NA* (94.9) | 99.9 (71.5) |
| **Refinement** |  |  |  |
| Reflections used in refinement | 24286 (2521) | 18673 (2898) | 17468 (1267) |
| Resolution range | 35.625 - 2.35<br>(2.44 - 2.35) | 29.83 - 2.50<br>(2.657 - 2.50) | 46.52 - 2.18<br>(2.24 - 2.18) |
| Reflections used for R-free | 1214 (133) | 898 (151) | 1746 (140) |
| Number of atoms | 4092 | 4039 | 2413 |
| *Macromolecules* | 3832 | 3757 | 2270 |
| *Ligands* | 90 | 80 | 49 |
| *Solvent* | 170 | 202 | 94 |
| Protein residues | 498 | 495 | 301 |
| B-factor | 40.83 | 37.5 | 51.4 |
| *macromolecules* | 41.0 | 37.6 | 51.68 |
| *ligands* | 40.3 | 33.9 | 43.26 |
| *solvent* | 38.1 | 37.0 | 48.88 |
| Ramachandran plot |  |  |  |
| *favoured* (%) | 96.94 | 96.92 | 98.33 |
| *allowed* (%) | 3.06 | 3.08 | 1.67 |
| Rotamer outliers (%) | 0.00 | 0.00 | 0.00 |
| **RMSD** |  |  |  |
| *bonds* (Å) | 0.004 | 0.0023 | 0.002 |
| *angles* (°) | 0.583 | 0.580 | 0.517 |
| Clashscore | 1.8 | 1.45 | 1.91 |
| $R_{work}$ | 0.1932 (0.2374) | 0.1906 (0.2309) | 0.205 (0.3048) |
| $R_{free}$ | 0.2318 (0.3217) | 0.2293 (0.2814) | 0.235 (0.3339) |

Statistics in parentheses are for the highest-resolution shell.
\* The HKL2000 version used for data processing did not give $CC_{1/2}$ value for overall data.

Interestingly, in one of the monomers, we could observe a chain of water molecules (bridging waters) that connects the attacking water to E66 through a series of hydrogen bonds (**Figure 1**). The interaction between the attacking water and E66 can also be represented through a different arrangement of bridging water molecules (**Figure S3**). This distant interaction involves a long chain of water molecules. B-factors for these waters are comparable to the average B-value for the main chain atoms of the protein – asserting the reliability of their positions in the active site. The significance of E66 in effectively positioning the attacking water is apparent from the structural analysis. Importantly, similar mechanisms for water-chain mediated nucleophile activation have been reported for both GTP and ATP hydrolyzing enzymes^15,49–51^, where a chain of water molecules mediates proton transfer from the attacking water to the respective catalytic residue(s). In addition to these insights, the structure of E67A mutant also provides a rationale for its slow catalytic rate. A comparison of all the protomers from *St*NFeoB E67A and *St*NFeoB WT structures reveals a significant variation in the arrangement of bridging waters along with variable conformations of E66 (**Figure S4A**). Therefore, we propose that this dynamic behavior of E66 and its extensive interaction with the chain of waters would decelerate the reaction rate in the E67A mutant, which is also evident from its *k*_cat_ of 0.89 min^-1^. In contrast, the other HAS-GTPases - MnmE and dynamin, show multi-fold higher catalytic rates of ∼50 min^-1^ and ∼200 min^-1^, respectively^15,16^ (**Figure S4C, S4D**). While, both these GTPases depict similar water-mediated catalysis, the water chains are much shorter, and the catalytic residues in both the cases are stabilized by additional interactions (**Figure S4C, S4D**). This stable arrangement would collectively help to reduce the entropic cost of aligning the catalytic machinery and could hence yield higher catalytic rates. All these analyses indicate towards a water-mediated nucleophile activation mechanism in *St*NFeoB E67A and underscore the importance of E66 for this reaction.

**Figure 1:**
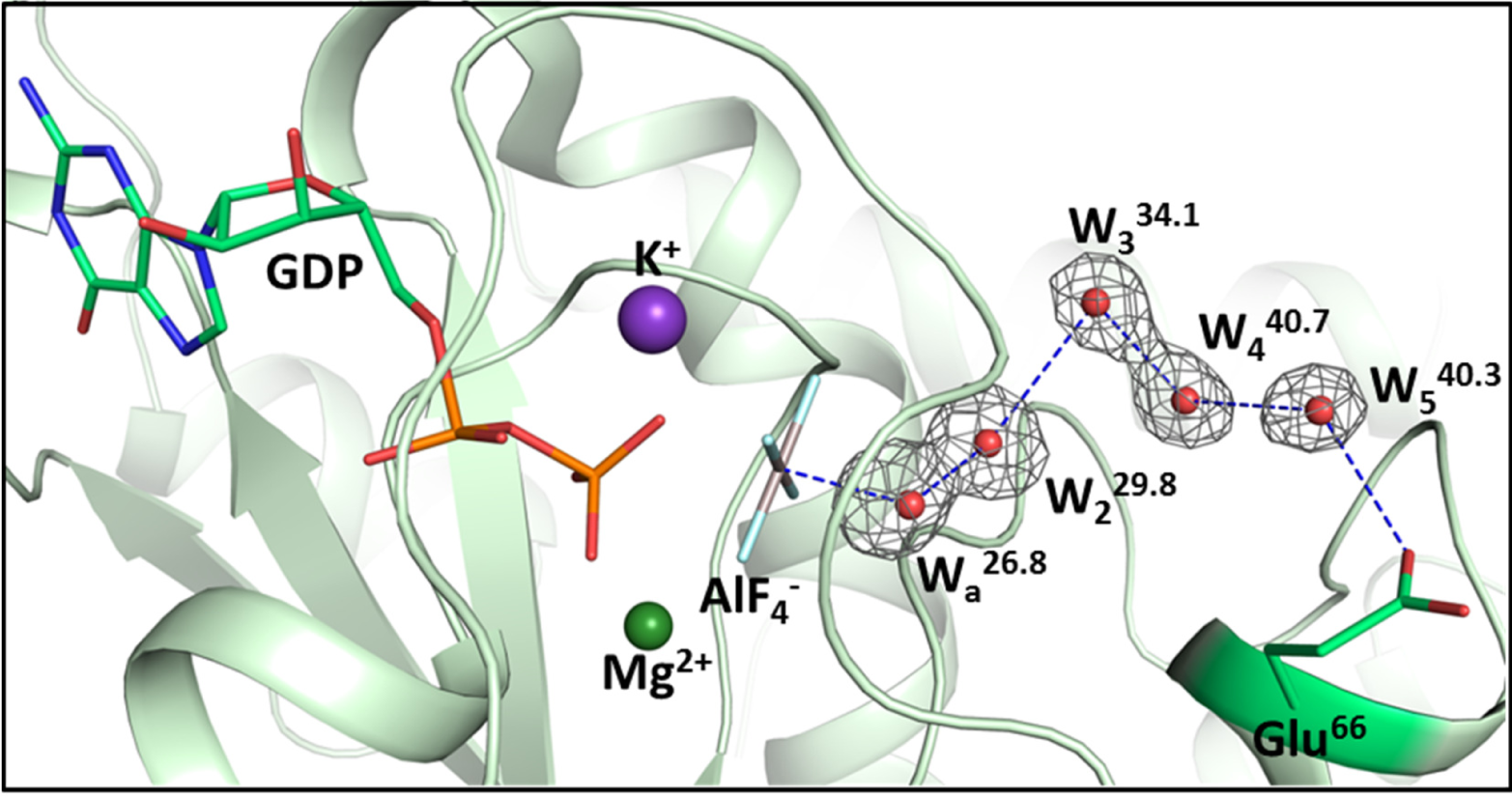
Active site arrangement of *St*NFeoB E67A. The transition state structure of *St*NFeoB E67A mutant (light green) illustrating the arrangement of active site waters and its interaction with Glu66 (green). The waters are labeled as W_a_ (attacking water) and W_2_-W_5_; the value in superscript denotes their respective B-factors. For comparison, the average B-value for the main chain atoms is 38.4. The grey mesh around the water molecules represents the Polder omit map contoured at 5σ. Hydrogen bonds are represented as blue dashes.

In accordance with the structural studies, previously performed QM/MM calculations further elucidate the role of E66^43^. E66 acts as the terminal proton acceptor and the proton from the nucleophilic water is transferred through a chain of bridging water molecules. Moreover, an analysis of different snapshots from the intermediate state of GTP hydrolysis simulations clearly highlights the dynamic nature of bridging water molecules and the catalytic residue (**Figure S4B**). However, the spatial organization of the active site waters and the catalytic E66 show distinct deviation with respect to that of the crystal structure (**Figure S4B**). All these structure models are a representative snapshot from an ensemble of possible conformations, and it may be difficult to precisely capture this transitory arrangement of waters in the crystal structures. Regardless of these variations, both the studies implicate the importance of E66 in positioning the attacking water through a chain of bridging waters. It can be concluded that in the absence of E67, residue E66 could play a catalytic role in *St*NFeoB. An important and unsettling concern however is the observation of GTP hydrolysis when the catalytic E66 is mutated to Ala^41^ (**Figure 2A**). To understand this, we attempted to crystallize the *St*NFeoB E66A mutant with GDP.AlF_x_ complex. Although, we managed to crystallize it in various conditions, we were unable to obtain single crystals despite exhaustive optimization trials. Therefore, we gather insights from our simulation studies^43^. These simulations indicated that in the absence of E66, residue E67 assumes the role of proton acceptor and drives catalysis in a similar manner. All these findings collectively suggest that in *St*NFeoB WT, either E66 or E67 can function as a proton acceptor. This also clarifies the observed GTP hydrolysis activity in each of the single mutants E66A and E67A (**Figure 2A**) and further rationalizes the strict conservation of this residue pair across FeoB homologues.

**Figure 2:**
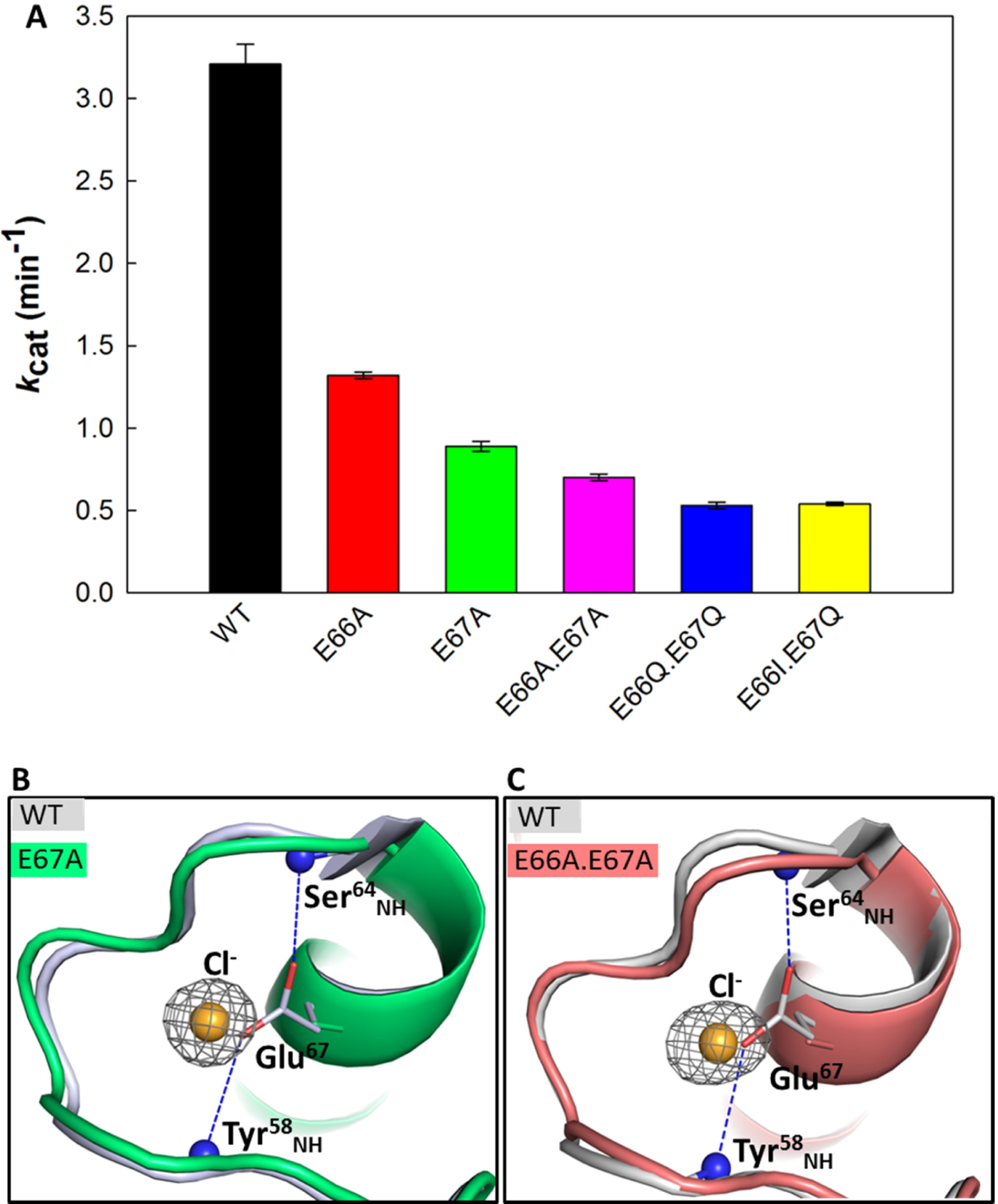
GTP hydrolysis of *St*NFeoB constructs. (A) The catalytic rates for GTP hydrolysis are determined through Michaelis-Menten kinetics. The values for WT, E66A, E67A and E66A.E67A have been taken from our previous study^43^. (**B**) and (**C**) Chloride ion occupies the position of the mutated residue, Glu67. The structural superposition of WT (grey) (PDB: 3SS8) with E67A (green) and E66A.E67A (salmon). The grey mesh represents the omit map (F_o_-F_c_) around the Cl^-^ (orange) contoured at 6σ in both the mutant structures. The interactions of E67 with its main chain in the WT structure are represented by blue dashes.

### StNFeoB continues to hydrolyze GTP in the absence of both Glu66 and Glu67

*St*NFeoB uses two mutually exclusive catalytic residues to perform the same function. We anticipated that mutation of both the glutamates would impair GTP hydrolysis. However, we were surprised to observe a considerable activity for the double mutant *St*NFeoB E66A.E67A, when compared to that of an internal control, *St*NFeoB T35A mutant (**Figure 2A** and **Figure S5**). T35 is a highly conserved residue in the switch-I loop that holds the critical cofactor Mg^2+^ and stabilizes the attacking water^2,4^. Mutation of this Thr impairs GTP binding as well as GTP hydrolysis. This unexplained GTP hydrolysis activity of the double mutant is incongruent with the catalytic role of the conserved Glu pair and necessitates further examination.

We determined the transition state structure of *St*NFeoB E66A.E67A to understand the structural basis of GTP hydrolysis (**Table 1** and **Supplementary information**). However, unlike in *St*NFeoB E67A, the crystal structure of double mutant could not identify a prospective residue that could act as a base, either by itself or through other interaction(s) in proximity (**Figures S6 and S7**). The water molecules mediating interactions between the attacking water and the protein, only involve the main chain atoms or polar residues (**Figure S6**). An extensive analysis of all the proximal polar residues (within 15 Å distance) to attacking water in search of a prospective catalytic residue, revealed only two residues, Y58 and S64, that are oriented towards the active site (**Figure S7**). However, either of these themselves cannot act as a base and in addition, are not coupled to a suitable/proximal proton acceptor. Moreover, mutating these residues did not affect GTP hydrolysis (in comparison to that of WT protein)^41^. We further examined the possible participation of the other monomer (of the asymmetric unit dimer) in triggering GTP hydrolysis. However, we could not find any plausible direct/indirect interactions from the other monomer either (**Figure S8A**). The residue located nearest from one monomer to the attacking water of the other is ∼22 Å away and there is no appreciable interaction between the two monomers. In conjunction, the analysis of surface packing for various crystallographic interfaces from PISA server^52^ and the elution profile of *St*NFeoB E66A.E67A (**Figure S8B**), also preclude the existence of a biologically relevant oligomeric structure for the double mutant. Therefore, despite thorough structural analysis, we could not assign a catalytic base for the double mutant.

We revisited our structures to evaluate another feature which we have overlooked so far, considering it to be a crystallization artefact. In both the structures of E67A and E66A.E67A mutants, we observed an unaccounted sphere of electron density (**Figure 2B, 2C**). Curiously, this unaccounted density coincides with the space previously occupied by the carboxyl moiety from E67. A similar density was also observed for a previously determined structure of *St*NFeoB E67A co-crystallized with GMPPNP (a non-hydrolyzable analogue of GTP)^40^. It was established that this density arose due to occurrence of a chloride ion that occupies the void created due to the substitution of E67 to Ala. Accordingly, we have also modeled a chloride ion in both the structures, as it aptly satisfies the density and also maintains the equivalent interactions with the protein backbone. Since chloride is an inherent component of ionic salts that are abundantly present in our crystallization solutions, and also as important constituents of the assay buffers. It is important to discern, if the occurrence of chloride, is a crystallographic artefact, or it holds any biochemical significance. Although, it is difficult to assign a catalytic role for chloride, its persistent occurrence in replacement of E67 necessitates a thorough investigation of its effect on catalysis in the double mutant.

To this end, we have generated a double glutamine mutant (*St*NFeoB E66Q.E67Q) since glutamines are thought to preserve the interactions analogous to glutamates, although they cannot themselves act as proton acceptors. Besides, the side chains of glutamines would occlude the binding of the chloride ion. However, upon characterization, the double glutamine mutant also exhibited GTPase activity equivalent to that of double alanine mutant (**Figure 2**). Similarly, another double mutant, E66I.E67Q was also investigated for its effect on GTP hydrolysis. We reasoned that this mutant, apart from occluding the chloride, would possess a bulky hydrophobic residue to interfere with the chain of waters that stabilize the attacking water. However, these mutations too have a minimal effect on the GTPase activity (**Figure 2**). Additionally, since none of the tested mutations impact nucleotide binding (**Table S1** and **Figure S9**), it is apparent that all the double mutants retain a catalytic potential and it is independent of the chloride ion in the active site.

### StNFeoB double mutant utilizes an alternative mechanism for GTP hydrolysis

The intriguing observation that GTP hydrolysis persists in the double mutants, made us to inquire the possible emergence of newer ‘alternative’ mechanisms for GTP hydrolysis, which may remain dormant otherwise, and emerge only when the catalytic glutamates are rendered inactive. However, despite thorough structural analysis we could not assign a catalytic base for the double mutant. It is likely that the double mutant utilizes an alternative catalytic mechanism which does not require a residue to serve as base. It should be noted that in contrast to *St*NFeoB WT or its single catalytic mutants, classical GTPases Ras and EF-Tu are known to employ substrate activated GTP hydrolysis. In this mechanism, the γ-phosphate of the substrate (GTP) deprotonates and activates the attacking water^23,24,32^. The molecular details of proton transfer cannot be inferred from the crystal structures using X-ray diffraction. Therefore, we resorted to computational methods involving hybrid quantum mechanical/molecular mechanical (QM/MM) metadynamics simulations, whereby we could elucidate the mechanism of GTP hydrolysis in the double mutant^43^.

The computational studies revealed that in the absence of catalytic glutamates, the arrangement of active site waters is altered significantly^43^. Despite this alteration in arrangement of water molecules the double mutant continues to hydrolyze GTP, albeit through a different means. The attacking water (W_a_) is now activated through proton transfer to the γ-phosphate of GTP, and importantly, an auxillary water molecule (W_aux_) plays a crucial role in this process by functioning as an intermediary (**Figure S10**). In addition, this catalytic machinery requires further reinforcement from accessory water molecules (W_3_ and W_4_), which stabilize a catalytically competent orientation of the water molecules (W_a_ and W_aux_). Interestingly, the arrangement of water molecules in monomer A of the double mutant crystal structure, presents a peculiar resemblance to intermediate snapshot from the metadynamics simulations (**Figure 3A**). Three of these active site water molecules from the crystal structure show a significant overlap with W_a_, W_aux_ and W_4_. Whereas, W_3,_ which bridges the W_aux_ and W_4_ in the transition state snapshot, do not align. Moreover, a relatively higher B-value suggests poor stability of this water. It is apparent from the snapshots obtained from the simulation that the position of W_3_ in the active site is contingent upon the precise molecular orientations of waters - W_aux_ and W_4_ (**Figure 3A**). We reason that this displacement of W_3_, in the crystal structure, may be due to an alteration in orientation(s) of active site water(s). Since, hydrogen atoms do not sufficiently scatter X-rays, the exact orientation of these water molecules cannot be inferred from these crystal structures. We speculate that these differences are a consequence of the inherent limitation associated with the use of AlF_4_^-^, which despite its efficient imitation of the planar configuration of phosphoryl intermediate, has a comparatively distinctive charge distribution. Notably, AlF_4_^-^ with its four equatorial fluorides (in contrast to the trigonal phosphoryl intermediate), along with the more electronegative nature of F^-^ and a large ionic radius of Al^3+^, render subtle, yet distinct, variations to its electrostatic configuration. These minute variations, although inconsequential for the bulky protein macromolecule, might have considerable implications for the precise molecular orientations and the arrangement of proximal water molecules. Nevertheless, the distinctive arrangement of active site waters can be easily appreciated from this transition state structure. Additionally, it allows meticulous analysis of the active site residues and investigation of their roles in GTP hydrolysis. Therefore, interpretations of these structural analyses, together with simulation studies clearly illustrate the conversion of catalytic mechanism in *St*NFeoB, from residue directed proton transfer in WT and single mutants to SAC in the double mutant.

**Figure 3:**
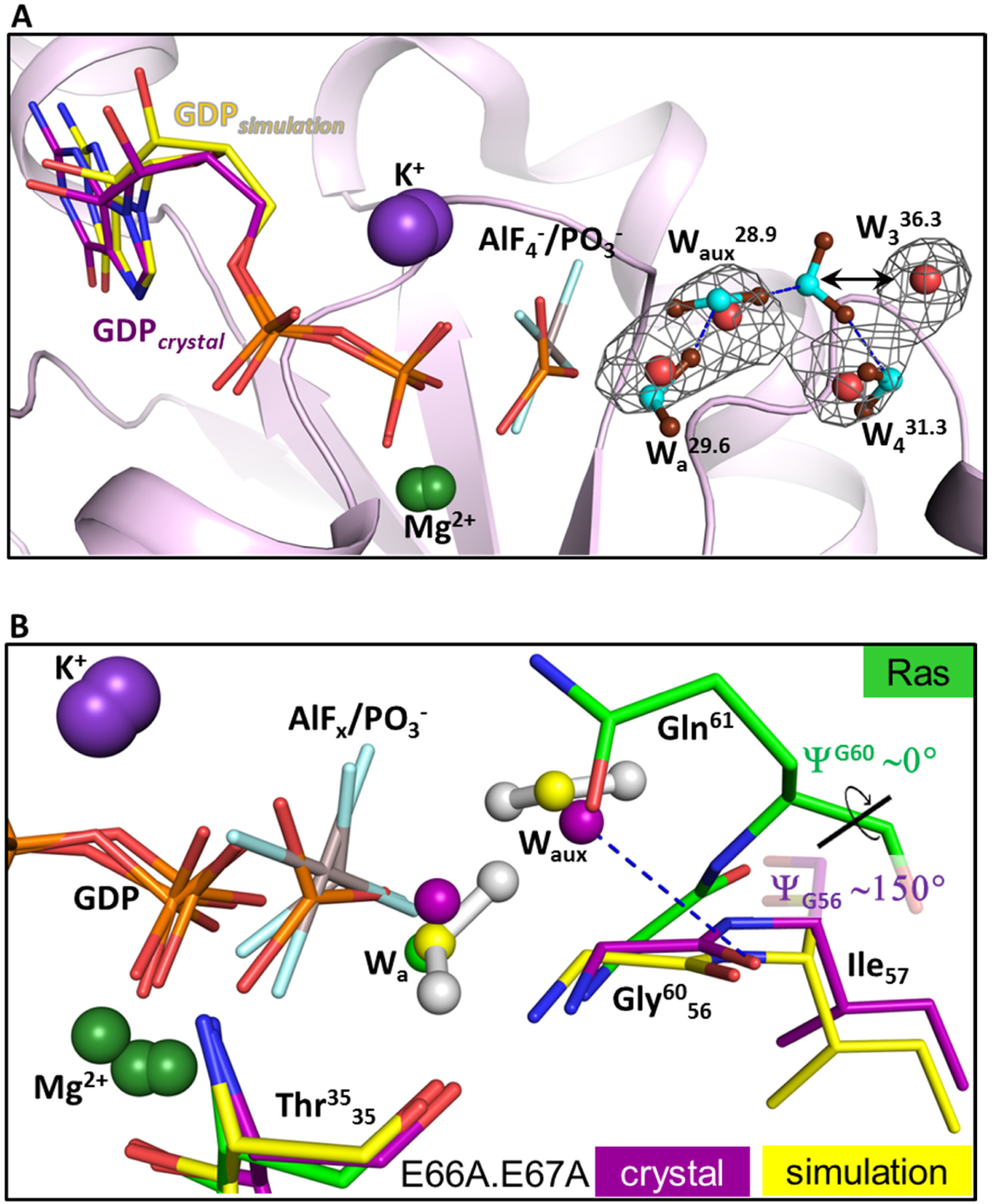
Active site arrangement of *St*NFeoB E66A.E67A. (**A**) The transition state structure (monomer A) of *St*NFeoB E66A.E67A mutant (light pink) illustrating the arrangement of active site waters. The waters are labelled as W_a_ (attacking water), W_aux_ (auxiliary water), W_3_ and W_4_; the value in superscript denotes their respective B-factors. For comparison, the average B-value for the monomer’s main chain atoms is 38.2. The grey mesh around the water molecules represents the Polder omit map contoured at 5σ. Crystal structure is superposed with a snapshot (yellow) from QM/MM simulations, representing the intermediate state of GTP hydrolysis^43^. The waters are shown as red spheres for the crystal structure and as ball and sticks – with cyan coloured oxygen and chocolate brown hydrogens – for the simulation snapshot. The hydrogen bond interactions between waters in simulation snapshot are shown through blue dashes. (**B**) Structural superposition of transition state structure of Ras (green) (PDB: 1WQ1), transition state structure of *St*NFeoB E66A.E67A (purple) and a snapshot from metadynamics simulations, representing the intermediate state of GTP hydrolysis in *St*NFeoB E66A.E67A (yellow). The waters are represented as spheres for crystal structures and as ball and sticks (with hydrogens) for the simulation snapshot. The oxygen atoms of waters are coloured in accordance with respective structure models. The numbers in residue labels represent numbering for Ras in superscript and for *St*NFeoB E66A.E67A in subscript.

Ras also utilizes a SAC mechanism for its GTPase activity; however, it is dependent on the catalytic Q61 which serves as an intermediary for proton transfer to the γ-phosphate^53^. It is curious to note that while a water molecule (W_aux_) functions similarly as Q61 in the double mutant of HAS-GTPase *St*NFeoB, a hydrophobic substitution of Q61 in Ras does not accommodate a water molecule and is rather deleterious for its function. It can be appreciated from structural analysis that W_aux_ occupies a similar position as the side chain of Q61 (**Figure 3B**). Therefore, the substitution of Q61 with a hydrophobic amino acid prevents the occurrence of a water molecule at this position. On the contrary, in *St*NFeoB or any other HAS-GTPase, the hydrophobic amino acid at the analogous position (I57) is retracted away from the active site to in turn create a void space at this position. When we first classified HAS-GTPases^33^, we realized that they harbor insertions in the switch-II region, which imparts distinct conformations to this loop and orients the hydrophobic residue away from the active center (**Figure 3B**). We further hypothesized that the void thus created, would present opportunities for the emergence of diverse catalytic mechanisms^33^. The observation of SAC in the double mutant provides yet another variation to the GTP hydrolysis mechanisms of HAS-GTPase. Moreover, since the features responsible for positioning W_a_ and W_aux_ are conserved across the HAS-GTPases^43^, we believe these findings have broader implications for GTP hydrolysis in HAS-GTPases. Therefore, HAS-GTPases might possess a basal catalytic activity, even in the absence of their catalytic residues.

Interestingly, a similar behavior was recently reported for another GTPase, hGBP1, where the pair, S73 and E99, was identified to play a catalytic role^51^. However, mutation of either of these residues, fails to abrogate GTP hydrolysis. The difference with regard to FeoB is that these mechanisms activate an alternative catalytic residue that serves as base. When S73 is mutated, two possibilities arise – a water molecule occupies the position of S73 and participates in water-chain mediated catalysis together with E99 or with D103. On the other hand, when E99 is mutated, D112 in its vicinity assumes the role of proton acceptor. However, it is yet to be established, if these mechanistic alternatives hold significance for the physiological function (i.e. Fe^2+^ transport in case of FeoB). Or, it is just a vestigial attribute, which became dormant during the course of evolution.

### Catalytic residue(s) in AaEra

Era is another HAS-GTPase, which functions in the assembly of 30S ribosomal subunit^44–46^. Like FeoB, it is also a cation-dependent GTPase, activity of which is stimulated in the presence of K^+ 54^ (**Figure S11B**). The insights gained from *St*NFeoB – its distantly located catalytic residues and long water-chain mediated catalytic mechanism – provide a perspective to understand GTP hydrolysis in Era. Furthermore, the alteration of GTP hydrolysis mechanism of *St*NFeoB in the absence of catalytic residues raises curious possibilities. Since the features that facilitate the SAC mechanism (in the double mutant of *St*NFeoB) are well preserved among all the HAS-GTPases^43^, we believe the catalytic mutants of HAS-GTPases would maintain biochemical activity through SAC mechanism. Therefore, to consolidate our understanding, we have studied the GTP hydrolysis in *Aquifex aeolicus* Era (*Aa*Era).

Structural superposition of *Aa*Era^55^ and a few HAS-GTPases^15,16,34,43^ with established catalytic residues helps to map their spatial organization with respect to *Aa*Era structure. Additionally, colouring the *Aa*Era structure based on the conservation of each residue across the Era homologues (using *Consurf* ^56^) is further expected to assist in the identification of prospective catalytic residue(s) (**Figure 4A**). The carboxylate moieties of catalytic residues E282 of MnmE and D152 of Atlastin occupy similar positions irrespective of their completely different spatial origins. Interestingly, the side chain of *Aa*Era Y63 is also positioned in proximity to the carboxylates of the above-mentioned residues in the superposed structures (**Figure 4A**). Moreover, this position is evolutionarily conserved among the Era homologues according to *Consurf* analysis, and no other conserved polar residue in the active site could be identified. However, Tyr cannot act as a catalytic base by itself; it has to be supported by an acidic residue. In the wt structure, Y63 of *Aa*Era interacts with D69, and could possibly serve the catalytic role for *Aa*Era (**Figure 4A**). Although distantly located, D69 could possibly act as an ultimate base, like the distant E66 or E67 in *St*NFeoB. The available structures of GMPPNP bound *Aa*Era, however, do not indicate the existence of a continuous chain of water molecules that connects the attacking water with Y63 (**Figure 4A**). These arrangements of catalytic machinery are transitory and may only be captured through their transition state structures; e.g., the catalytically competent conformation of the conserved Gln and its apparent role in G_α_ and Ras proteins was not evident until their transition state structures were determined^11–13^.

**Figure 4:**
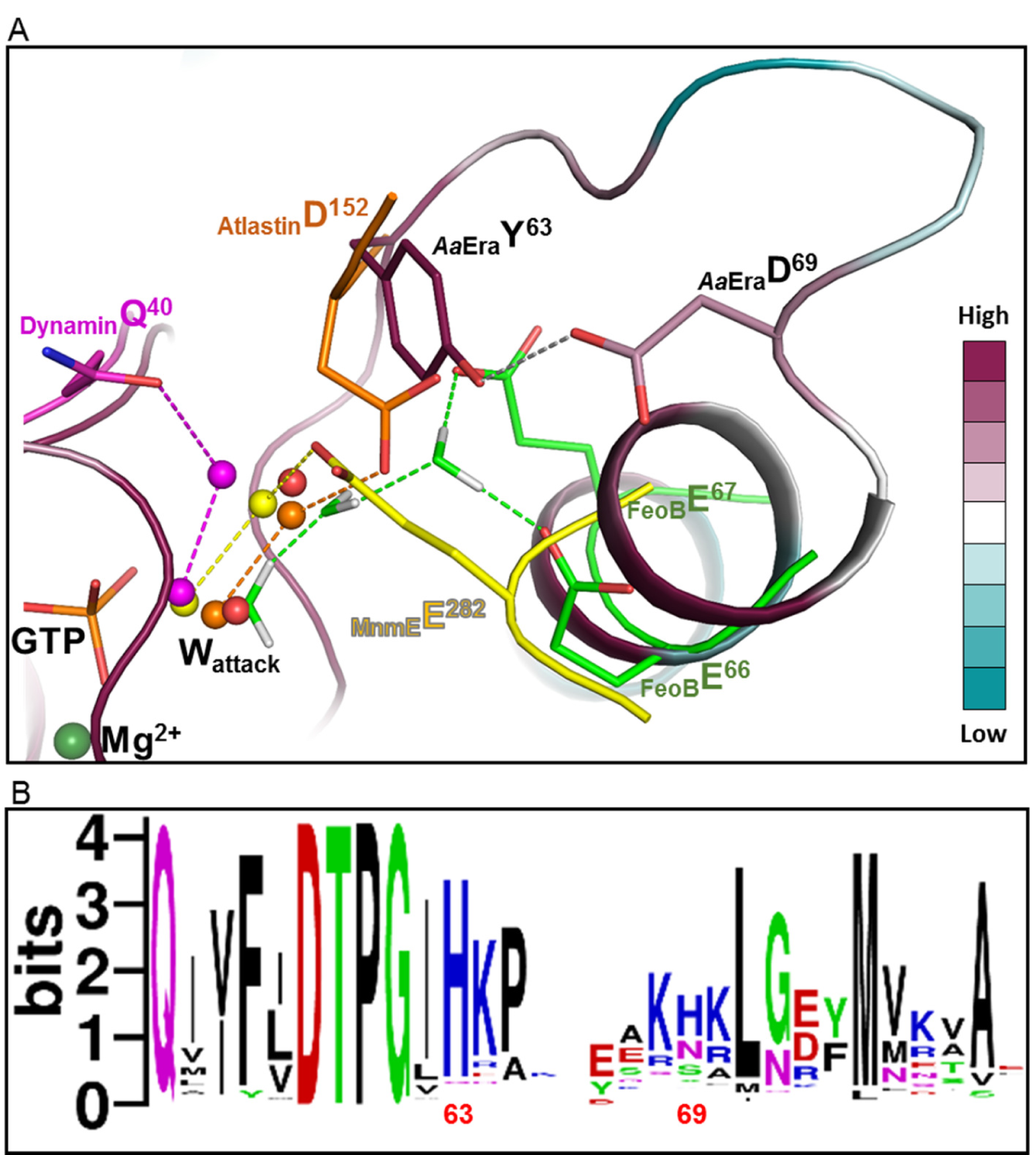
Potential catalytic residue(s) in *Aa*Era. **(A)** Active site architecture of *Aa*Era (PDB: 3IEV), coloured (maroon to turquoise) on the basis of conservation score for each residue through *Consurf. Aa*Era is superposed with structures of other HAS-GTPases – Atlastin (orange) (PDB: 4IDO), Dynamin (pink) (PDB: 2×2F) and MnmE (yellow) (2GJ8) – bound to transition state analogue (GDP.AlF_x_); and simulation snapshot of transition state of GTP hydrolysis in FeoB^43^ (green). Only the catalytic residue(s) and active site waters (in respective colours of their structures) are shown for structures other than *Aa*Era. Active site waters in the *Aa*Era structure are shown as red spheres. The hydrogen bond interactions of active site waters and respective catalytic residues are shown through dashed lines and coloured accordingly. **(B)** Sequence logo representation (generated using *WebLogo*) of the aligned sequences of Era homologues used in *Consurf* analysis. Positions corresponding to residues 63 and 69 of *Aa*Era are labeled.

Intriguingly, in spite of significant conservation scores for positions 63 and 69 (**Figure 4A)**, the residues Y63 and D69 are not well represented among the Era homologues (**Figure 4B**). Tyr seems to be inexistent and is rather substituted by a highly conserved His, while D69 is mostly substituted by a polar residue (**Figure 4B**). Additionally, it is located in a loop region which has a variable length due to insertions or deletions among the homologues. Although the position corresponding to residue 63 of *Aa*Era is conserved owing to the occurrence of His, further substitutions – although infrequent – could be observed in the homologues analyzed through *Consurf* (**Figure S12A**). This inadequate representation of Era homologues with Tyr and other infrequent substitutions at the position corresponding to residue 63, prompted us to examine a larger set of Era sequences. Using *PSI-BLAST* ^57^, we have retrieved and analyzed sequences from distantly related homologues. We have used ∼17000 sequences for analysis, and surprisingly, only 14 of these homologues have Tyr at the position corresponding to residue 63 of *Aa*Era (**Figure S12B**). 9 of these 14 homologues belong to the phylum *Aquificae* of extreme thermophilic bacteria. All these nine homologues possess a conserved Y63-D69 pair; although, not all the *Aquificae* phylum homologues have Y63 (**Figure S13**). Despite variations in position 63, it is interesting to note that an acidic residue is conserved at position corresponding to residue 69 or (in some cases) adjacent to it (**Figure S13**). These organisms from *Aquificae* phylum belong to extreme niches and are hence poorly represented in this sequence analysis. Nevertheless, this small subset of Era sequences is quite diverse, showing variations up to 50 %.

The sequence analysis suggests that the combination of a polar and an acidic residue could have significance for GTP hydrolysis in Era homologues from the *Aquificae* phylum. This is further corroborated by a significant decrease in the biochemical activity of D69 mutants (**Table 2, Figure 5** and **Figure S14**). Mutation of D69 results in a ∼5-fold decrease in GTP hydrolysis, which, although significant, cannot be regarded as an abrogation in the activity. Earlier, we have elucidated that mutation of both the catalytic residues E66 and E67 in *St*NFeoB activates SAC mechanism for GTP hydrolysis. Therefore, if D69 serves as the catalytic base, its mutation may activate the SAC mechanism to continue hydrolyzing GTP. Alternatively, the mutation of D69 might activate some other residue to compensate for its loss. Such a diversion in mechanism has been reported for hGBP1^51^. Intriguingly, mutation of Y63 does not reduce GTP hydrolysis; rather, there is an increase (∼2-fold) in catalytic rate (**Figure 5 and Table 2**). It is curious to note that this persistence of biochemical activity in Y63 mutants resembles the catalytic mutant S73A of hGBP1. The mutation of S73A maintains its catalytic activity by using a chain of water molecules to mediate transfer of proton to either E99 (canonical base) or to D103 (alternative base, which activates upon mutation of S73A). The sequence and biochemical analyses suggest the catalytic importance of the Y63-D69 pair in *Aa*Era. Simultaneously, it poses questions regarding the possibilities of hydrolytic activities of the mutants.

**Table 2:** Michaelis-Menten kinetic parameters for *Aa*Era proteins

| | $v_{\max}$ | $K_M$ |
| --- | --- | --- |
| *AaEra* Y63A | $28.36 \pm 1.06$ | $850.53 \pm 92.12$ |
| *AaEra* WT | $15.48 \pm 0.41$ | $755.47 \pm 61.93$ |
| *AaEra* D69A | $3.40 \pm 0.06$ | $356.60 \pm 28.82$ |

**Figure 5:**
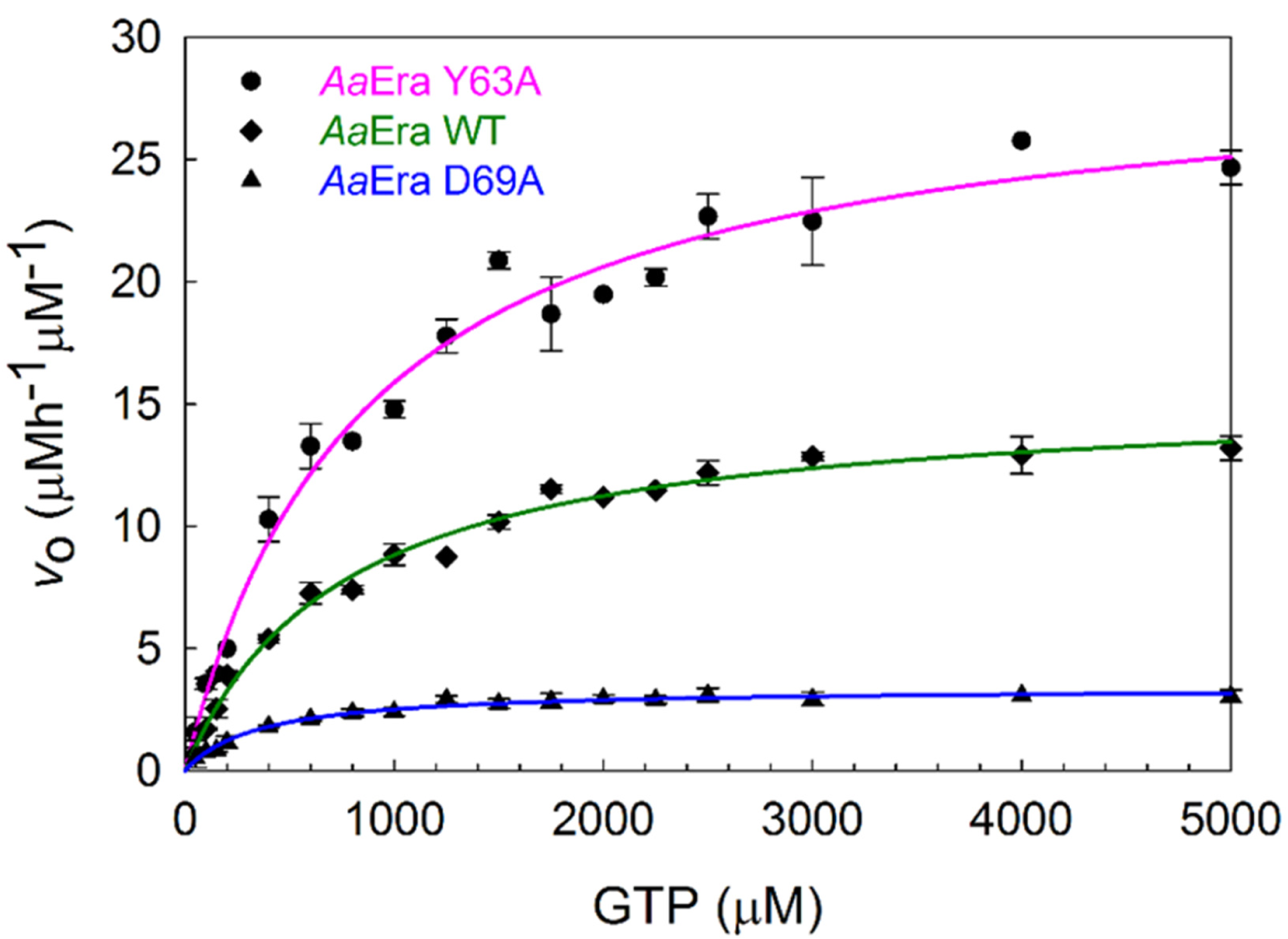
GTP hydrolysis activities of *Aa*Era constructs. Michaelis-Menten kinetics for *Aa*Era constructs.

### AaEra utilizes substrate assisted catalytic mechanism for GTP hysrolysis

Crystallization trials with *Aa*Era wt did not yield any crystals. Therefore, we gather insights from QM/MM simulations to understand its catalytic mechanism (Neha Vithani, Ph.D. Thesis^58^). Contrary to the above propositions regarding involvement of Y63-D69 pair in GTP hydrolysis, these simulations do not indicate any interactions between the attacking water and residue Y63, and instead reveal a SAC mechanism for GTP hydrolysis in *Aa*Era (**Figure 6A and Figure S15B**). Similar to the *St*NFeoB double mutant, the attacking water (W_a_) in *Aa*Era wt is also activated by proton transfer to the terminal phosphate. This proton transfer is mediated by the auxiliary water molecule (W_aux_). Despite uniform conservation of this feature among all the HAS-GTPases, SAC remains dormant in FeoB wt and is activated only upon mutation of both the catalytic glutamates. The structure analysis of *Aa*Era wt active site reveals the distinct features that impose SAC as the default catalytic mechanism. In the wt structure^55^ (PDB: 3IEV), Y63 and M75 pack closely to form *van der Waals* interactions, resulting in a hydrophobic surface that prevents access to the hydroxyl group of Y63 (**Figure 6B**). This hydrophobic barrier hinders the interaction of attacking water with distantly located D69.

**Figure 6:**
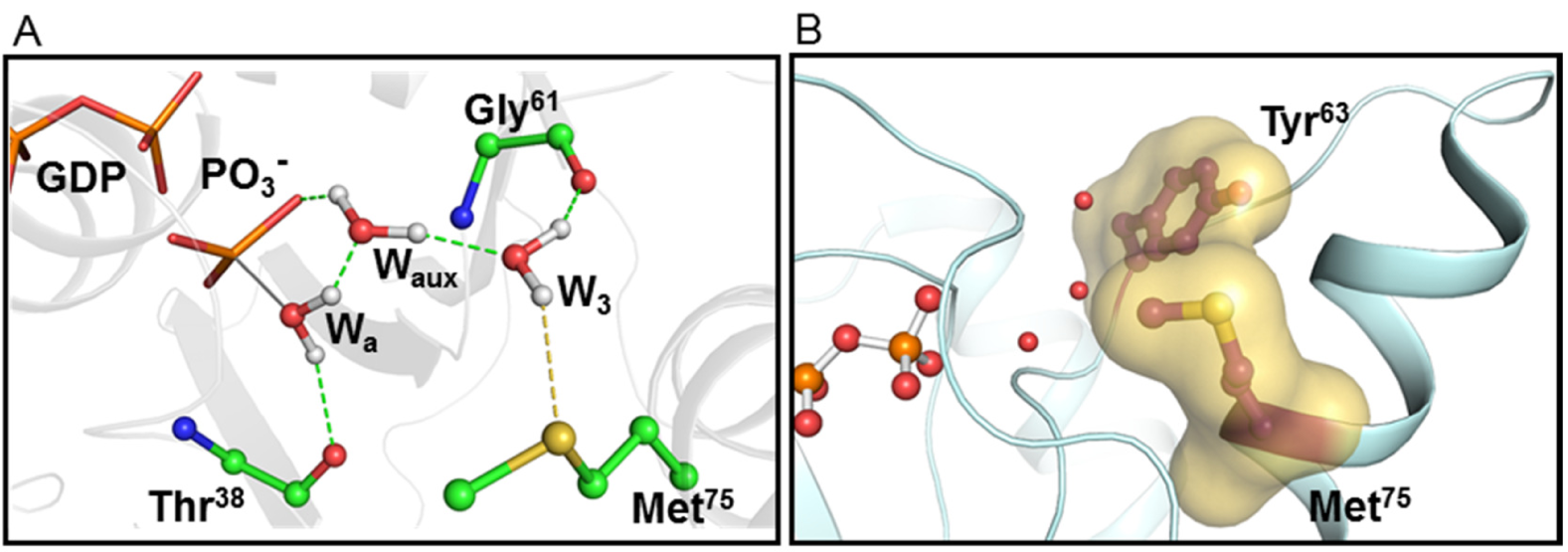
GTP hydrolysis in *Aa*Era WT. **(A)** Transition state snapshot of GTP hydrolysis from QM/MM simulations on *Aa*Era WT (Neha Vithani, Ph.D. thesis, IIT Kanpur^58^). The arrangement of active site waters (shown in ball and stick representation) with hydrogen bonds shown as green dashes. Polar interaction with M75 is represented as a yellow dash and the bond formation of attacking water through a solid grey line. **(B)** Surface representing the interface between Y63 and M75 in the active state structure of *Aa*Era WT (PDB: 3IEV).

M75 plays another important role in initiating GTP hydrolysis. Simulations reveal that M75 changes from an ‘inward’ facing conformation in the GTP-bound reactant state to an ‘outward’ facing conformation of catalytically active state (**Figure S15A**). This alteration in the conformation of M75 allows the re-orientation of active site waters, which enables them to transfer a proton from the nucleophilic water to the γ-phosphate (**Figure 6A** and **Figure S15**). The relevance of these minute distinctions in the orientation of water molecules can be further appreciated through observations from FeoB^43^. In the presence of either E66 or E67, an extended chain of waters is formed with their hydrogen atoms pointing towards glutamate(s). However, mutation of both the glutamates releases this restrain on waters, allowing them to re-orient and thus favor the SAC mechanism (**Figure S10**). Similarly, in *Aa*Era, M75 regulates GTP hydrolysis by applying constrains on the orientation of active site waters. Importantly, even when the methyl group of M75 flips out, the hydroxyl group of Y63 side-chain remains inaccessible to active site waters (**Figure S16A**). Conclusively, it is the interface between Y63 and M75 that enforces and regulates the SAC mechanism of GTP hydrolysis in *Aa*Era wt.

The position corresponding to M75 is highly conserved among the Era homologues, and is occupied by a Met or Leu (**Figure S16B**). On the other hand, the position corresponding to residue 63, which forms the interface with residue at position 75, is rarely occupied by a Tyr. Since the interface between Y63-M75 governs the SAC mechanism in *Aa*Era, the variations at position 63 might have implications for the catalytic mechanism of Era homologues. A significant number of Era homologues harbor a bulky hydrophobic residue Phe (13 %) or Ile (6 %) at this position (**Figure S16B**). The occurrence of bulky hydrophobic residues suggests the formation of a strong hydrophobic interface with M75, which is expected to seclude the active site waters and impose SAC mechanism for GTP hydrolysis among these homologues. However, in a majority of homologues the position corresponding to Y63 is substituted by a polar His (67 %) and Gln/Asn (8.5 %). A lack of conserved acidic residue in the vicinity of residue 63, among these homologues, indicates towards the existence of SAC mechanism. Histidine can itself function as a proton acceptor depending upon its microenvironment; therefore, a side-chain mediated mechanism of nucleophile activation cannot be completely ruled out. There is an ambiguity over the involvement of His in catalysis, and a comprehensive structural and biochemical analysis of different homologues would be required to understand the unification and diversity of catalytic mechanisms among Era homologues. Nevertheless, the simulation and structural analysis on *Aa*Era wt indicates the importance of Y63-M75 interface for GTP hydrolysis. It is intriguing that mutation of Y63 increases GTPase activity rather than impeding it. This warrants further investigation to understand the reason for efficient GTP hydrolysis in the Y63 mutants.

### Alternative mechanism of GTP hydrolysis in AaEra Y63A

We have determined the crystal structure of *Aa*Era Y63A, in the presence of GDP.AlF_x_ complex, to understand the underlying catalytic mechanism. These crystals diffracted to a resolution of 2.18 Å and belonged to the space group P2_1_2_1_2_1_. The Y63A structure has been refined to R_work_/R_free_ of 0.205/0.235. Relevant statistics are listed in **Table 1**. The final structure models all the residues (1-301) of the protein. The nucleotide-binding site of the Y63A structure further contains the transition state analog, GDP and aluminum tetra-fluoride complex (GDP.AlF_4_^-^), a magnesium ion (Mg^2+^); and importantly, the structure provides evidence for binding of a potassium ion (K^+^) in the active site (**Figure S17**) (see **Supplementary information**).

This active state structure of Y63A allows us to observe the molecular details of its active site. Analysis of the hydrogen bonding network of the nucleophilic water indicates that it interacts with distantly located D69 through a series of four intermediary water molecules (**Figure 7A**). However, this long chain of water molecules depicts strong intermolecular hydrogen bonds with an average acceptor-donor pair (O-O) distance of 2.8 Å. Moreover, the B-factors for these waters are comparable to the average B-value for main chain atoms, and further augment the reliability in their position (**Figure 7A**). The molecular architecture of the active site suggests that in the absence of Y63 residue, D69 could be sufficient to activate the attacking water using a chain of connecting/bridging waters. In the crystal structure of *St*NFeoB E67A, we observed a similar five water-mediated catalytic arrangement. This long chain of waters, however, has a decelerating effect on GTP hydrolysis, which is also evident from the slow catalytic rate of Y63A (*k*_cat_ = 0.49 min^-1^). Interestingly, this activity is comparable to that of 0.89 min^-1^ for mechanistically related *St*NFeoB E67A.

**Figure 7:**
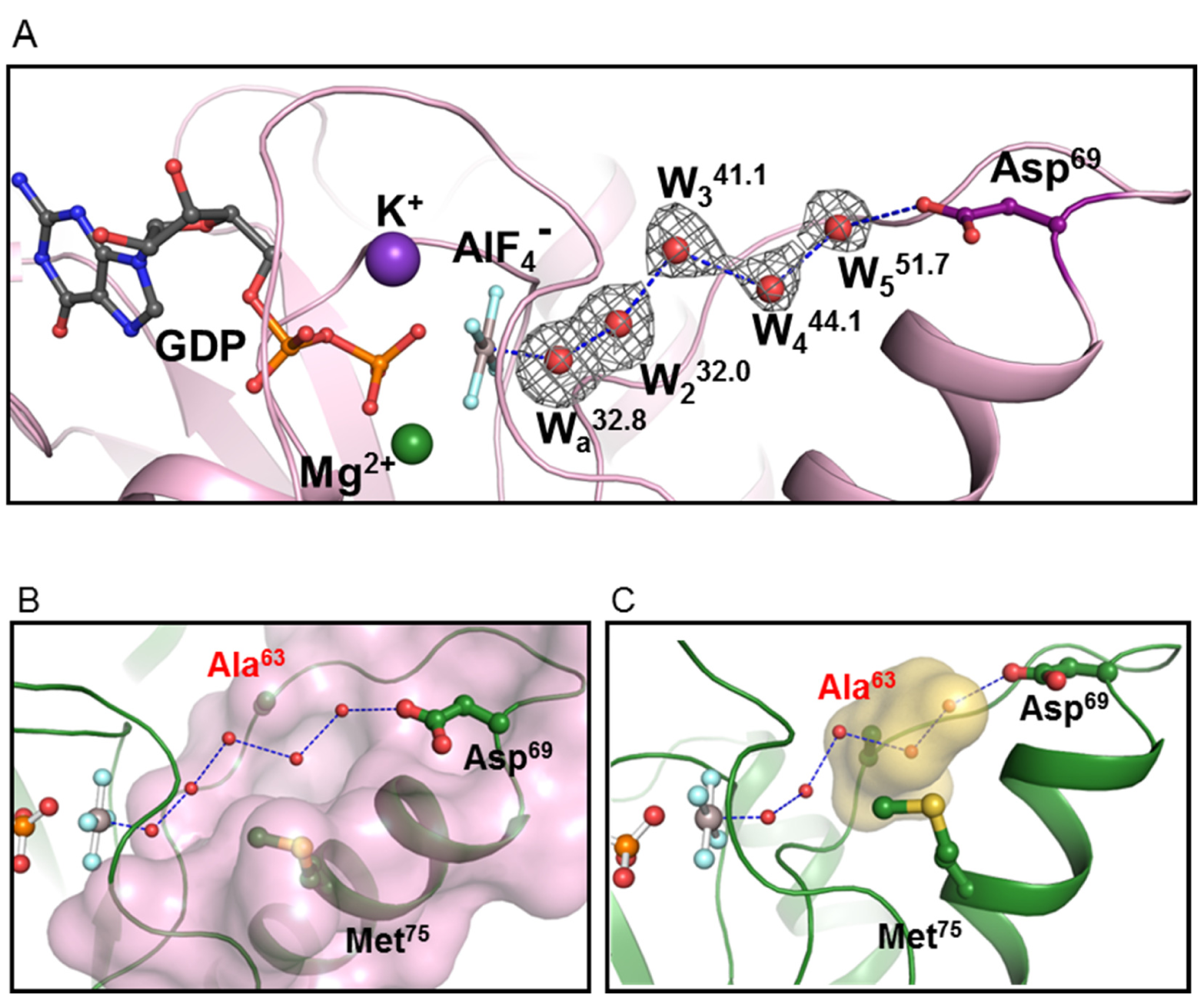
Active site arrangement of *Aa*Era Y63A. **(A)** The active site of the *Aa*Era Y63A mutant illustrating the arrangement of active site waters and its interaction with D69 (magenta). The waters are labeled as W_a_ (attacking water) and W2-W_5_. The value in superscript denotes their respective B-factors. The average B-value for the protein main chain is 51.3. The grey mesh around the water molecules represents the Polder omit map contoured at 5σ. **(B)** Surface representation of the active site of *Aa*Era Y63A in pink colour. **(C)** Arrangement of active site waters in the structure of *Aa*Era Y63A mutant; previously occupied space by Y63 in WT structure is represented as yellow surface

As noted earlier, the interface between Y63 and M75 of *Aa*Era wt hampers the interaction of attacking water with D69. Interestingly, the mutation of Y63 to Ala creates a conduit that allows the chain of waters to interact with D69 directly and opens an alternative route for activation of GTP hydrolysis in *Aa*Era (**Figure 7B & 7C**). While the wt utilizes SAC mechanism for GTP hydrolysis; it seems the Y63A mutant activates the nucleophile by proton transfer to D69, similar to the glutamates of *St*NFeoB wt.

### GTP hydrolysis in EcEra

The interface between the residues 63 and 75 seems to be crucial for regulating GTP hydrolysis among Era homologues. Therefore, we have investigated the significance of this interface in catalysis of *Ec*Era. *Ec*Era has a H67 at the position corresponding to Y63 of *Aa*Era and shows a slower rate of GTP hydrolysis in comparison to *Aa*Era (**Figure S11B**). Interestingly, mutating the H67 to bulky hydrophobic residues, Phe or Trp, does not impact GTPase activity (**Figure 8**). These bulky residues are thought to maintain the interface with M78. The observation of equivalent activities in wt and H67F or H67W suggests that H67 might play a structural role rather than a direct catalytic role – analogous to Y63 of *Aa*Era. Interestingly, the mutation of H67A shows a marked increase in hydrolysis, similar to the Y63A mutant of *Aa*Era. This increase in activity indicates either a transition to a side chain mediated catalysis, believed to be functional in *Aa*Era Y63A, or a structural re-arrangement of the active site that increases the efficiency of SAC. While it is difficult to specify the reason for the increase in activity without further structural and computational analysis, it is tempting to speculate that either of the glutamates E69-E70 – that follow H67 in *Ec*Era – could act as a proton acceptor in the absence of H67. Furthermore, we attempted to deliberately alter the catalytic mechanism through a substitution of H67D, which we expected to function as a strong base. However, this mutant surprisingly shows a significant reduction in activity (**Figure 8**), attributed primarily to a significant reduction in GTP binding (**Figure S18D**). The proximity among three strongly acidic side chains (D67, E69 and E70) might be responsible for the destabilization of the active site and poor nucleotide binding.

**Figure 8:**
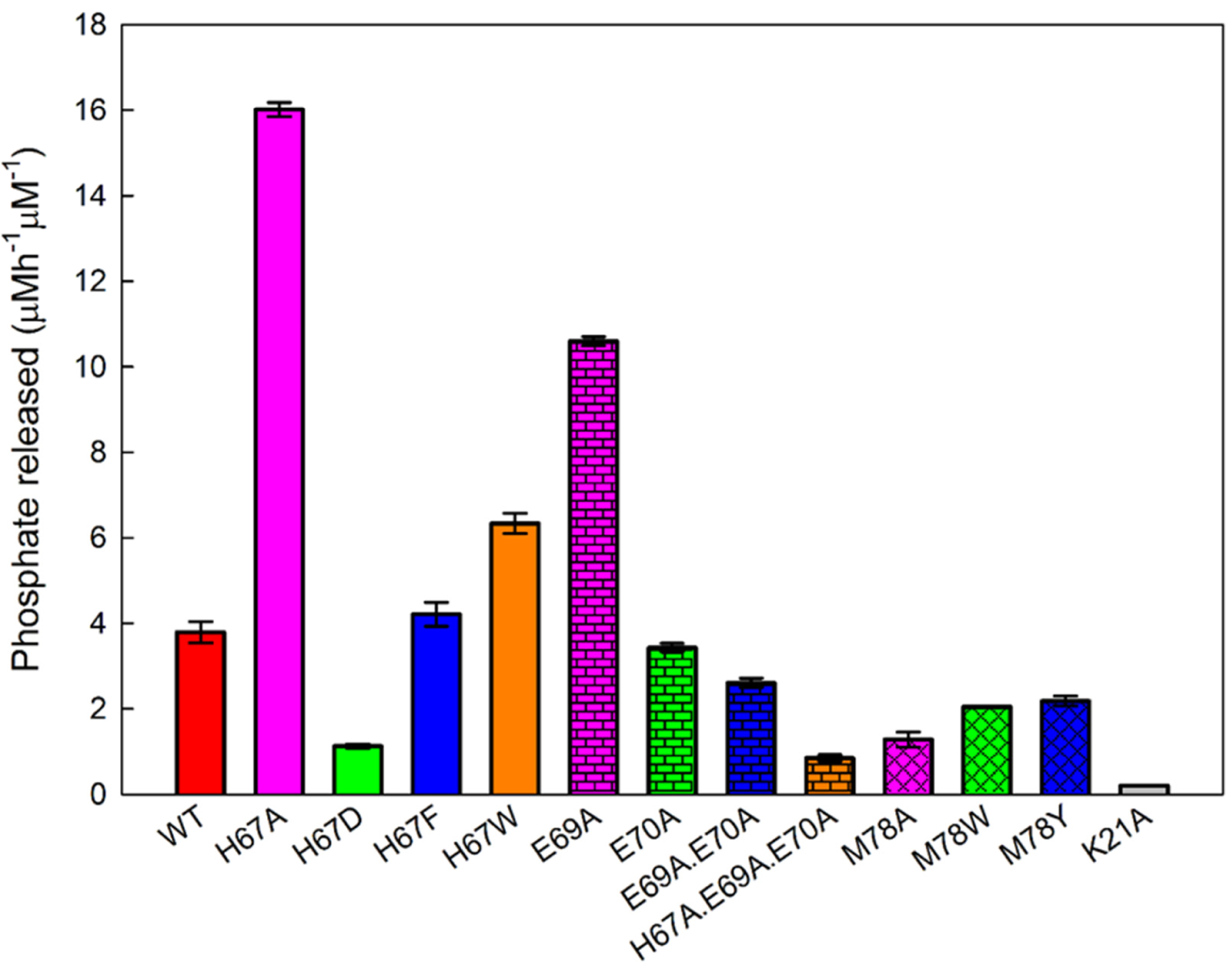
GTP hydrolysis of *Ec*Era constructs. Comparison of GTP hydrolytic activities of different *Ec*Era constructs.

The plausible involvement of E69 and E70 in the alteration of GTP hydrolysis by these mutants prompted us to investigate their significance for *Ec*Era. Mutation of both E69 and E70 shows little effect on the activity (**Figure 8**), indicating that primarily the interface between H67-M78 governs GTP hydrolysis of *Ec*Era and is not affected by substitutions in the switch-II region. However, one of the single mutants E69A again shows a significant increase in GTP hydrolysis (**Figure 8**). This sudden increase in hydrolytic activity highlights the conformational flexibility of the switch-II loop and alludes towards the transformation of the catalytic mechanism, as observed in the H67A mutant. Therefore, we decided to mutate all the three residues to construct *Ec*Era H67A.E69A.E70A (Triple mutant). Interestingly, this triple mutant shows a marked reduction in hydrolysis and truly represents a compromised catalytic state (**Figure 8**), as unlike H67D, it binds GTP efficiently (**Figure S18E**). Moreover, the activity of the triple mutant is drastically reduced in comparison to the H67A mutant, suggesting an important catalytic role for either E69 or E70 in the H67A mutant. Further comparing the activities of the double mutant E69A.E70A with the triple mutant re-emphasizes the relevance of interface between H67 and M78 in the effective organization of active site waters. The significance of this interface and importance of M78 for the formation of this interface is further evident through the characterization of M78A mutant, which shows poor nucleotide binding owing to the destabilizing effect it has on switch-II (**Figure S18F**), as observed from MD simulations (Neha Vithani, personal communication). Additionally, the substitution of M78 with bulky hydrophobic groups Trp and Tyr also show mildly compromised hydrolytic activities (**Figure 8**). Perhaps, the rigid side chains of these substituted residues are able to maintain the interface with H67, although, they do not possess flexibility, like the side chain of Met. Nevertheless, with all these mutants of *Ec*Era, we have generated a suite of constructs that depict GTP hydrolysis to variable extents that 1) span an order of magnitude (H67A/E69A *vs*. H67D/M78A/Triple mutant), 2) can be attributed to different molecular bases, i.e. binding and hydrolysis (H67D/M78A *vs*. Triple mutant) and 3) possibly employ different catalytic mechanisms (H67A/E69A *vs*. wt). In summary, this analysis of *Ec*Era mutants points towards the significance of the interface between H67 and M78 for GTP hydrolysis. In addition, these modulations in activities of interface mutants highlight the flexibility of *Ec*Era, which retains its catalytic ability – apparently, by activating alternative routes for nucleophile activation.

## Summary and outlook

This work addresses the emergence of alternative catalytic mechanisms in two HAS-GTPases, Era and FeoB. Here, we present a combination of structural and biochemical investigations to elucidate these mechanisms. In FeoB, we observe a pair of conserved glutamates, located distantly from the γ-phosphate, that perform the catalytic function. These residues utilize an extended chain of water molecules for proton transfer and initiate GTP hydrolysis. Interestingly, mutation of these catalytic residues invokes a re-arrangement of active site water molecules, and activates an alternative mechanism for GTP hydrolysis, by allowing the γ-phosphate to serve as the ultimate proton acceptor^43^. This terminal phosphate mediated activation of attacking water is known as substrate assisted catalytic (SAC) mechanism. Intriguingly, Era utilizes the SAC mechanism for GTP hydrolysis and exclusively depends on the arrangement of active site waters^58^. The investigation of Era homologues, presented here, could not identify an exclusive residue(s) that would function as a catalytic base. However, we observed the occurrence of a conserved interface at the active site, which constrains the water molecules and promotes the SAC mechanism. Surprisingly, disruption of this interface still maintains GTP hydrolysis in both *Aa*Era and *Ec*Era homologues. The disruption of the interface allows the active site residues to access these water molecules, which seem to activate side-chain mediated ‘alternative mechanisms’ for GTP hydrolysis. These ‘alternative mechanisms’ as observed in Era and FeoB, provide robustness against the mutation of the so-called catalytic residues. Similar emergence of alternative mechanisms was reported in hGBP1^51^. Taken together, this work allows us to question the notion of a catalytic residue; we hypothesize that persistent GTP hydrolysis seems indispensable for such systems to function based on the biological context.

HAS-GTPases have hydrophobic residues at the position corresponding to Gln^cat^ of Ras. The substituted hydrophobic residues show a retracted conformation and were envisaged to utilize distantly located catalytic residues (either from other locations in G-domain or from other interacting partners)^33,59^. It is apparent from the structural analysis that these HAS-GTPases supply the catalytic residue from diverse three dimensional positions^15,16,34^. The catalytic residues from these HAS-GTPases interact with the attacking water through a bridging water molecule and compensate for their distant origin^59^. We now learn that unique structural features, shared by all the HAS-GTPases, stabilize this bridging water molecule^43^. In FeoB, the bridging water is part of the water chain that serves as an intermediary for proton transfer to either of the catalytic glutamates. However, this bridging water reorients itself in the absence of catalytic glutamates, and continues to function as an auxiliary water molecule that mediates proton transfer to the terminal phosphate^43^. While this auxiliary water provides structural plasticity to FeoB, it is the basis for catalysis in Era. It is intriguing how these GTPases tailor their mechanisms according to the biological context. Most of the HAS-GTPases are universally conserved among the prokaryotes and are considered to be the ancestral GTPases^60^. Therefore, the SAC mechanism, as observed in Era and the double mutant of FeoB, could be the ‘primitive’ mechanism in the progenitor HAS-GTPases.

Another important observation from both of these HAS-GTPases is their dependence on an extensive arrangement of active site waters for GTP hydrolysis. Incidentally, mutations of catalytically important residues, in the active site of either of these, lead to an altered arrangement of active site waters. This flexibility seems to confer structural plasticity and provides robustness to the molecular switches. It is possible that the alternative mechanisms, as observed here, are the safeguard-mechanisms that maintain the functionality of these molecular switches. An extension of this work that comprehensively analyzes structure-function relationships among several HAS-GTPases would be necessary to establish the importance of the emergence of alternative mechanisms reported here.

## Acknowledgements

We are grateful to Dr. Neha Vithani and Prof. Nisanth Nair, IIT Kanpur for scientific inputs and sharing the simulation snapshots for preparing illustrations. We are thankful to Dr. Simon Goto and Prof. Hyouta Himeno, Hirosaki University for kind gift of *E. coli* Era expression construct; and for stimulating discussions. We thankfully acknowledge Prof. David Waugh, National Cancer Institute, NIH for the generous gift of *Aquifex* Era expression construct. We are thankful to Prof. Bichitra Biswal, Mr. Ravi Kant Pal and X-ray crystallography facility, National Institute of Immunology, New Delhi for data collection of *St*NFeoB crystals. We are also thankful to X-ray facility, National Centre for Biological Sciences, Bangalore for data collection of *Aa*Era Y63A crystals. We thankfully acknowledge contributions from all the previous and present members of BP lab; in particular, thanks to Dr. Pratik R. Patil for critical discussions and inputs on this work. We also thank AK lab members for support. BP thanks Department of Science and Technology for funding this work through grant DST/JSPS/P-248/2017. AK acknowledges Department of Science and Technology for funds through DST/NM/NT-2018/48.

## Author Contributions

SB and BP conceived the study and planned experiments. SB performed the experiments under BP’s supervision. AK contributed resources. SB and BP analyzed the results and wrote the manuscript.

## Methods

### Construction of expression plasmids

#### *St*NFeoB expression plasmids

The DNA encoding NFeoB domain (*St*NFeoB) that corresponds to residues 1-270 was amplified using genomic DNA from *Streptococcus thermophilus* (ATCC) as a template and cloned using NdeI and HindIII in the modified pGEX expression vector^61^, that bears a TEV protease cleavage site to remove the N-terminal GST tag from the fusion protein. All the mutants of *St*NFeoB were generated through the overlap PCR method, using the pGEX-*St*NFeoB WT vector as a template. The primers (**Table S2**) for specific substitutions (with codon sequence(s) underlined) were used to amplify the plasmids and circularized using the in-house Gibson assembly mix^62^ and transformed in *E. coli* DH_5_α cells. All the mutations were confirmed through DNA sequencing.

#### *Aa*Era expression plasmids

The expression plasmid for *Aa*Era (pBA1971) was a generous gift by Prof. David Waugh, NIH. The plasmid has been used to express *Aa*Era WT protein for crystallization experiments^55,63^. The plasmid expresses *Aa*Era along with N-terminal MBP-tag, which can be removed using TEV protease. The TEV cleavage site is followed by a 6XHis-tag, followed by sequence coding for *Aa*Era. All the constructs for expression of *Aa*Era mutants were generated through an overlap PCR method, using the pBA1971 as template. The primers have been listed in **Table S3**. Underlined sequences represent the mutant codons for specific substitutions. The plasmids were amplified using these primer pairs, and amplicons were circularized using Gibson assembly mix and transformed in E. *coli DH*5α cells. All the constructs were confirmed through DNA sequencing.

#### *Ec*Era expression plasmid

The expression plasmid for *Ec*Era (pTXera) was a generous gift by Dr. Simon Goto and Prof. Hyouta Himeno, Hirosaki University. The plasmid expresses an un-tagged *Ec*Era protein. All the constructs for expression of *Ec*Era mutants were generated similarly, as mentioned above. The primers are listed in **Table S4**.

### Protein expression and purification

#### *St*NFeoB proteins

The *St*NFeoB proteins were expressed and purified as reported previously^41^. Briefly, the cell lysates containing GST tagged protein(s) in Buffer A: 20 mM Tris (pH-7.6) and 100 mM NaCl, were loaded on GSTrap HP (5 ml) cartridge (GE Healthcare). The bound protein(s) were treated with TEV protease to release the untagged *St*NFeoB proteins from the column. These were subsequently purified through gel permeation chromatography on Superdex 75 10/300, or Superdex 200 PG 26/600 column (GE Healthcare). It was pre-equilibrated in Buffer B: 20 mM Tris (pH-8.0) and 100 mM NaCl. The fractions containing proteins were concentrated and stored at -80 °C after flash-freezing.

#### *Aa*Era proteins

*Aa*Era proteins were expressed in *E. coli* Rosetta (DE3) cells. The cells were induced at 30 °C in the Auto-induce media^64^. The cells were harvested and processed for protein purification, according to previously established protocol^55^. Briefly, the cell suspension in Buffer C: 50 mM Tris (pH -7.2), 200 mM NaCl and 1 mM Benzamidine, was lysed. The clarified lysate was treated overnight with TEV protease at room temperature. After TEV treatment, NaCl was increased to 1 M final concentration, and the lysate was heat-treated at 80 °C for 30 min to precipitate most of the host proteins. The clarified lysate was then loaded on HisTrap HP (5 ml) cartridge (GE Healthcare), and eluted through a linear gradient of Buffer D: 50 mM Tris (pH -7.2), 1 M NaCl and 500 mM Imidazole. These proteins were subsequently purified through gel permeation chromatography on Superdex 200 column (GE Healthcare) in Buffer E: 20 mM Tris (pH-7.2) and 150 mM NaCl. The fractions containing proteins were concentrated and stored at -80 °C after flash-freezing.

#### *Ec*Era proteins

*Ec*Era proteins were expressed in *E. coli* BL21(DE3) cells. The cultures were induced at 18 °C in the Auto-induce media^64^ and processed for protein purification post-harvesting. The cells were lysed in Buffer F: 20 mM Tris (pH-7.6), 10 mM KCl and 1 mM PMSF. The lysate was loaded on a HiTrap Q FF (5 ml) cartridge (GE Healthcare), pre-equilibrated with the same buffer. The bound *Ec*Era was eluted through a linear increment of Buffer G: 20 mM Tris (pH-7.6), 150 mM KCl. Fractions containing *Ec*Era protein were combined and diluted to reduce the salt concentration, and loaded on pre-packed HeparinHC-650M beads (Tosoh Biosciences), equilibrated with Buffer F. The protein was eluted by applying a linear gradient of Buffer H: 20 mM Tris (pH-7.6), 300 mM KCl. The relevant fractions were concentrated and stored at -80 °C after flash-freezing.

### GTPase activity assays

*St*NFeoB proteins (2-5 µM each) were incubated with a range of substrate (GTP for molecular biology, Thermo Scientific) concentrations (25 µM to 5000 µM), to determine the kinetic parameters (*k*_cat_ and K_M_). The reactions were performed at 37 °C, in GTPase assay buffer, Buffer R: 20 mM Tris (pH-8.0), 200 mM KCl and 5 mM MgCl_2_. The reactions were quantified through Biomol Green reagent (Enzo Life Sciences), a malachite green-based phosphate detection system. The rate of GTP hydrolysis (phosphate release) was plotted for each substrate concentration and the values were fit to rectangular hyperbolic function for deriving the kinetic parameters. Similarly, *Aa*Era proteins (1-7.5 µM each) were incubated with a range of substrate concentrations (50 µM to 5000 µM) to determine initial velocities at each concentration.

For comparing the GTPase activities of different mutants, the proteins were incubated with 500 µM GTP at 37 °C for 1 h. The reactions were terminated by adding Biomol Green reagent and the colouration developed after 20 min of stopping the reaction is measured through by taking the absorption at 620 nm and quantified using a phosphate standard.

### Crystallization

#### *St*NFeoB proteins

The proteins were diluted in Buffer R: 20 mM Tris (pH-8.0), 200 mM KCl and 5 mM MgCl_2_, and incubated with 1 mM GDP, 1 mM AlCl_3_ and 10 mM NaF on ice for 2-3 h. *St*NFeoB E66A.E67A mutant was diluted to a final concentration of 7.5 mg/ml. The protein-ligand mixture was mixed in 1:1 ratio with mother liquor: 100 mM Tris (pH-8.0), 400 mM NH_4_Cl and 26 % (w/v) PEG 6000. Similarly, *St*NFeoB E67A mutant was diluted to 7 mg/ml and mixed in 1:1 ratio with mother liquor: 100 mM Tris (pH-8.0), 50 mM NH_4_Cl and 30 % (w/v) PEG 6000. The crystals were obtained through vapor diffusion in the sitting drop setup at 18 °C.

#### *Aa*Era Y63A

The protein was diluted to achieve a concentration of 10 mg/ml in GTPase activity buffer, Buffer R. The protein was incubated in the presence of 2 mM GDP, 2 mM AlCl_3_ and 20 mM NaF at 72 °C for 15 min and allowed to cool down, slowly. The protein mix was then centrifuged at room temperature to pellet minute precipitate generated during the heating step. This protein-ligand mixture was mixed in equal amount with the mother liquor: 100 mM Bis-Tris (pH-7.6), 200 mM Ammonium acetate and 18 % (w/v) PEG 10,000. The crystals were obtained through vapor diffusion in a sitting drop setup at 18 °C.

### Data collection, processing and structure determination

#### *St*NFeoB crystals

The crystals of StNFeoB E67A and StNFeoB E66A.E67A both grew under a high concentration of PEG 6000, and hence, they were harvested and mounted directly on the goniometer. The crystals were diffracted using X-rays of wavelength 1.54 Å, generated through Rigaku FR-E+ SuperBright rotating-anode generator. The data was collected at 100 K on a Rigaku R-axis IV^++^ detector. The data for StNFeoB E67A was processed using the *XDS* program^65^. *Phaser* ^66^ was used for the determination of initial phases using GDP.AlF_4_^-^ bound StNFeoB WT structure^41^ (PDB: 3SS8) as the intial structure for molecular replacement (MR). The final structural model was generated through an iterative process of manual model building in *Coot* ^67^ and structure refinement using *Phenix*^68^.

The data for StNFeoB E66A.E67A was processed using the HKL2000 program^69^. Again, MR was used for generating initial structure models using the same WT structure^41^ (PDB: 3SS8) as the model. MR was performed using online web-server *Auto-rickshaw*^70,71^. The initial model was improved through further rounds of manual model building using *Coot*^67^ and refinement. The structure refinement was performed using *Refmac5*.*5*^72^ and *Phenix*^68^. The Polder maps were generated in *Phenix*^73^. The representative graphic illustrations from the structures were prepared using *PyMOL* (Schrödinger, LLC) and *UCSF Chimera*^74^.

#### *Aa*Era Y63A crystal

The crystals were harvested and skimmed through mother liquor supplemented with 15 % (v/v) MPD to provide cryo-protection and mounted on the goniometer. The crystals were diffracted using X-rays of wavelength 1.54 Å, which were generated through rotating anode, Rigaku FR-X generator. The diffraction was performed at 100 K and Rigaku R-axis IV^++^ detector was used for data collection. The data was processed through *XDS* program^65^ and initial phases were determined through MR using *Phaser*^66^, with the active state structure of *Aa*Era WT bound to GMPPNP^55^ (PDB: 3IEV) as search model. The final structure was built through a series of refinement and structure building rounds in *Phenix*^68^ and *Coot*^67^, respectively. The omit maps were generated using Polder maps in *Phenix*^73^. LLG based anomalous difference map was constructed using *Phaser*, using the final structure model and un-merged reflection data as inputs for SAD experimental phasing. The graphic illustrations and structural analysis were prepared using *PyMOL* (Schrödinger, LLC) and *UCSF Chimera*^74^.

### Fluorescent nucleotide binding assays

Guanosine nucleotides labeled with fluorescent N-methyl-anthranoyl (mant) groups O-linked with 2’ or 3’ of ribose (Jena Biosciences), were used for binding studies. 5 µM of *St*NFeoB proteins were incubated on ice for 5 min with 0.5 µM of mant-GDP or mant-GMPPNP (a non-hydrolyzable analogues of GTP) in Buffer R supplemented with 1 mM DTT, which contains 20 mM Tris (pH-8.0), 200 mM KCl, 5 mM MgCl_2_ and 1 mM DTT. Similarly, 5 µM of *Aa*Era proteins and 10 µM of *Ec*Era proteins, each, were incubated briefly on ice with 0.25 µM of mant-nucleotide in Buffer R, supplemented with 1 mM DTT, prepared and added fresh. The fluorescence spectrum was measured on Perkin Elmer LS55 fluorimeter. The samples were excited using the light of wavelength 340 nm (slit width = 10 nm) and the emission from 400 nm to 600 nm (slit width =10 nm) was recorded.

### Sequence analysis

Initial analysis of conservation at each position was performed using the *Consurf* web-server^56^. Subsequently, a larger set of sequences for sequence alignment of Era homologues was retrieved using *PSI-BLAST*^57^. Homologues with incomplete sequences or those with extra domains were excluded from further analysis. Sequences with more than 90 % identity were removed using *CD-HIT*^75^, and remaining sequences were aligned using the *Clustal Omega*^76^ multiple sequence alignment tool from EBI web-server. The sequence alignments were analyzed using *SeaView*^77^ and *ALINE*^78^ was used to generate representative alignments for illustrations. Sequence logo representations of sequence alignments were generated from the WebLogo server^79^.

## Notes

### Competing Interest Statement

The authors have declared no competing interest.

