## Supplementary information for "Alternative catalytic mechanisms driven by structural plasticity is an emerging theme in HAS-GTPases, Era and FeoB"

*Structures of StNFeoB mutants represent an active state of GTP hydrolysis*

We have determined the crystal structures of mutants, *St*NFeoB E67A and *St*NFeoB E66A.E67A, co-crystallized in the presence of GDP and aluminium fluoride (GDP.$AlF_4^-$ complex). Both the mutants crystallized in similar conditions. The crystals of *St*NFeoB E67A and *St*NFeoB E66A.E67A diffracted to a resolution of 2.35 Å and 2.5 Å, respectively and belonged to space group $P2_12_12_1$ with similar unit cell dimensions. The final structure of *St*NFeoB E67A was refined to $R_{work}/R_{free}$ values of 0.1932/0.2318, and that of *St*NFeoB E66A.E67A was refined to $R_{work}/R_{free}$ values of 0.1906/0.2293. The statistics and details of data processing and refinement are listed in **Table 1**. The crystal structures depict similar packing of macromolecules, with two monomers in the asymmetric unit. However, there are minute differences between the two chains. In chain A of *St*NFeoB E67A structure, residues 1-256 could be modeled, and there is lack of electron density for residues 162-167; whereas, chain B has residues 1-255 and lacks density for residues 161-168. Similarly, in *St*NFeoB E66A.E67A structure, chain A has residues 1-255, lacking the density for residues 162-166; and chain B has residues 1-254 with insufficient density for residues 160-168. The lack in density for residues in these regions might be due to close contacts with adjacent monomers from other asymmetric units. Nevertheless, the root mean square deviation (*rmsd*) over all the common $C_\alpha$ atoms between the two chains of each structure is 0.35 Å for *St*NFeoB E67A and 0.25 Å for *St*NFeoB E66A.E67A. Therefore, both the chains can be treated equally for structural analysis. Importantly, both structures represent the conserved domain architecture and preserved inter-domain interactions between the G-domain (residues 1-160) and the adjacent helical domain (residues 173-255) of NFeoB (**Figure S2A**).

The nucleotide binding site of both the mutants contains a GDP.$AlF_4^-$ complex; a magnesium ion ($Mg^{2+}$) and a potassium ion ($K^+$) (**Figure S2B, S2C**). GDP.$AlF_4^-$ complex acts as a transition state mimic of GTP hydrolysis and can potentially shed light on the reaction mechanism[1–3]. Therefore, to understand the molecular details of catalysis in the mutants of *St*NFeoB, we determined the structures of these mutants in the active state of GTP hydrolysis and compared them with the previously determined active state structure of *St*NFeoB wildtype (WT)[4]. This comparison yields crucial information that allows us to decipher details of the reaction mechanism.

A strong structural similarity between the crystal structures of the mutants (*St*NFeoB E67A and *St*NFeoB E66A.E67A) and the active state structure of *St*NFeoB WT[4] (PDB ID: 3SS8), may be noted based on the overall *rmsd* of 0.63 Å and 0.59 Å, respectively. More importantly, the conformations of switch regions (Switch-I & Switch-II) in the mutant structures show a strong overlap with those of the WT structure (**Figure S2A**). Switch regions undergo conformational changes upon nucleotide hydrolysis[4–7]. Hence, their conformations as well as the conservation of characteristic interactions between the signature motifs (G2 and G3 motifs) and the active site ligands are considered to be important indicators of the active state structure. Switch-I or the G2 motif is best represented by a conserved threonine[5,8] (Thr35 in *St*NFeoB); the side chain provides coordination to the $Mg^{2+}$ and its main chain stabilizes the attacking water molecule (**Figure S2D**). Similarly, the switch-II region is characterized by the presence of conserved DxxG motif (G3 motif)[5,8]. While the glycine of this motif (Gly56) also makes interaction with the attacking water (**Figure S2D**), the conserved aspartate residue (Asp53) helps in stabilizing the $Mg^{2+}$ ion by positioning its coordinating water molecule. Analysis of switch loops from both the structures of mutants clearly highlights these fundamental interactions and indicates their active state conformation.

The importance of ordered switch regions to stabilize the $Mg^{2+}$ at the active site, underscores its indispensable role in GTP hydrolysis. $Mg^{2+}$ draws the electrons away from the γ-phosphate, resulting in the development of a partial positive charge that stimulates the nucleophilic attack by water molecule[9]. Other than the switch-I Thr35, $Mg^{2+}$ is hexa-coordinated to Thr15; oxygens of β-phosphate and γ-phosphate (in these structures, $F^-$ of $AlF_4^-$) and two water molecules (**Figure S2D**). While a $Mg^{2+}$ at active site is critical for catalysis, it alone is incapable to accelerate the hydrolysis reaction. The nucleophilic attack of water leads to the generation of strong negatively charged intermediate; which needs to be stabilized. It has been shown that *St*NFeoB uses a $K^+$ for this function[4,6]. Both the structures of mutants maintain the $K^+$ and its characteristic interactions *viz* the carbonyl oxygens of Gly29 and Trp31 from switch-I loop and side chain oxygen of Asn11 from the P-loop. Additional interactions are provided by the phosphates of the nucleotide (**Figure S2E**). Conclusively, all these observations suggest that both the structures indeed represent the active state of GTP hydrolysis. Therefore, the molecular details deduced from these would be reliable to interpret the catalytic mechanisms in the *St*NFeoB mutants.

*Crystal structure of AaEra Y63A*

GDP.$AlF_x$ serves as a mimic for an intermediate state of GTP hydrolysis[2]; here, $AlF_x$ represents a mixture of aluminum trifluoride ($AlF_3$) and aluminum tetrafluoride ($AlF_4^-$), that are generated *in situ* through the reaction of aluminum chloride ($AlCl_3$) and sodium fluoride (NaF).[10,11] The electron density from the omit map of the Y63A crystal structure clearly indicates the occurrence of $AlF_4^-$, which occupies the site of γ-phosphate and is also stabilized by the analogous interactions, i.e. the backbone amides of- G36, T37 and T38 from switch-I, and G61 from switch-II and N13 from P-loop (**Figure S17**). $AlF_4^-$ wedges between the β-phosphate of the nucleotide and the attacking water, with the aluminum atom forming, co-ordinate bonds of 2.1 Å and 2.2 Å with respective oxygen atoms F**igure S17A**). Therefore, the precise placement of this negatively charged planar moiety at short distances from the attacking and scissile groups makes $AlF_4^-$ an efficient mimic of γ-phosphate undergoing hydrolysis.

GTP hydrolysis causes the re-distribution of one electron equivalent of charge from γ-phosphate to β-phosphate[12] and hence, depends on positively charged moieties to accelerate hydrolysis[13]. GTP hydrolysis commences through an attack of nucleophilic water on the γ-phosphate and utilizes a conserved P-loop lysine and a $Mg^{2+}$ to initiate hydrolysis. It is suggested that both of these work in concert to draw the electron cloud away from γ-phosphate, which results in the induction of partial positive charge and hence, the nucleophilic attack[9,12]. In the current structure, K16 and $Mg^{2+}$ are positioned on the opposite sides of GTP, and each makes interactions with fluorine from $AlF_4^-$ and non-bridging oxygen from β-phosphate in their respective vicinity (**Figure S17A & 17C**). In addition, $Mg^{2+}$ forms coordinate bonds with S17 from the G1 motif, T38 from the G2 motif and two water molecules to complete its coordination sphere (**Figure S17A**). Apart from $Mg^{2+}$ and P-loop lysine, another positively charged moiety is presented in the form of either 'Arg finger'[5,14], or monovalent ions[13,15] to accelerate GTP hydrolysis. Insertion of this moiety into the active site drives the triphosphate chain of GTP towards an eclipsed conformation, which induces repulsion among the oxygen atoms of the phosphates and eventually favors bond cleavage[13,16,17].

Structural analysis suggests that Era possesses conserved features to hold a $K^+$ in the active site (**Figure S11A**), and this is corroborated by acceleration in GTP hydrolysis in its presence (**Figure S11B**). However, since previous crystallization experiments did not include $K^+$, a structural evidence for this does not exist.[18,19] Based on the reasoning above, we included $K^+$ in the buffer used for crystallization. Interestingly, a strong electron density in the $F_o$-$F_c$ omit map for Y63A structure suggests the occurrence of an ion at the active site. Moreover, an analysis of the likelihood-based anomalous difference map further indicates the existence of an anomalous scatterer at this position (**Figure S17B**). Therefore, we have modeled a $K^+$ in this density as 1) it appropriately satisfies the electron density; 2) overlaps with the position of $K^+$ in other $K^+$ dependent GTPases: NFeoB[6] and MnmE[20]; 3) makes ionic interactions with long bond lengths (averaging 2.83 Å)[21], characteristic of $K^+$ and 4) in addition, no regular geometry is associated with these interactions, which precludes the existence of any transition metal at the binding site. Therefore, this provides structural evidence for the significance of $K^+$ in GTP hydrolysis by Era. $K^+$ interacts with an oxygen atom from each of α-phosphate and β-phosphate and fluorine from $AlF_4^-$. It is stabilized by the backbone carbonyls of S32 and K34 from switch-I and side chain of highly conserved N13 from the P-loop (**Figure S17A**). Therefore, mutation of the homologous Asn in *Ec*Era abolishes the accelerating effect of $K^+$ on GTP hydrolysis[22] and further justifies the placement of $K^+$ in the active site of the Y63A structure. The structure further rationalizes the inadequacy of $Na^+$ in accelerating GTP hydrolysis of Era (**Figure S11B**). $Na^+$ has stronger charge density and forms relatively shorter ionic bonds (~2.4 Å) as compared to $K^+$ [21], thus limiting its interactions with the aforementioned residues. This lack of stabilization would restrict the binding of $Na^+$ to the cation binding site of Era. It is informative to contrast this observation with dynamin that utilizes $Na^+$ for stimulation of its activity[23]. The cation binding site of dynamin has been modified suitably to efficiently utilize $Na^+$ [13,15]. Overall, it can be concluded that the structure of Y63A preserves all the crucial features associated with the active state of GTP hydrolysis. Furthermore, the structure shows excellent superposition with GMPPNP bound wt structure with an *rmsd* of 0.5 Å over all the common $C_\alpha$ atoms, indicating once again that the Y63A structure represents the active state.

**Table S1: Michaelis-Menten constants for *St*NFeoB proteins.**

| ***St*NFeoB** | **$K_M$** |
|---|---|
| WT | 373 ± 69 μM |
| E66A | 276 ± 37 μM |
| E67A | 477 ± 68 μM |
| E66A.E67A | 314 ± 51 μM |
| E66Q.E67Q | 187 ± 20 μM |
| E66I.E67Q | 229 ± 21 μM |

**Table S2: Primers for performing polymerase chain reactions.**

| **Primer** | **Sequence (5' to 3')** |
|---|---|
| WT FP | GGAATTCCATATGACAGAAATTGCATTA |
| WT RP | CCCAAGCTTACGACAAGGTTAAAG |
| E66A FP | CTACAGTCCTGCCGAAAAGGTGG |
| E66A RP | CCACCTTTTCGGCAGGACTGTAG |
| E67A FP | GTCCTGAAGCCAAGGTGGCG |
| E67A RP | CGCCACCTTGGCTTCAGGAC |
| E66A.E67A FP | CTACAGTCCTGCCGCAAAGGTGGCG |
| E66A.E67A RP | CGCCACCTTTGCGGCAGGACTGTAG |
| E66Q.E67Q FP | CAGTCCTCAACAAAAGGTGGCGCGTGATTATTTG |
| E66Q.E67Q RP | ACCTTTTGTTGAGGACTGTAGGGTGACATTGAG |
| E66I.E67Q FP | CAGTCCTATACAAAAGGTG |
| E66I.E67Q RP | CCTTTTGTATAGGACTGTAG |
| T35A FP | GGTTGCTGTTGAGCGAAAGAGTGGTC |
| T35A RP | CGCTCAACAGCAACCCCTGGCCAG |

**Table S3: Primers for mutagenesis of *Aa*Era expression vector.**

| Primer | Sequence (5’ to 3’) |
|---|---|
| Y63A FP | CGGTATAGCCGAGCCGAAAAAG |
| Y63A RP | CGGCTCGGCTATACCGGGTG |
| Y63F FP | CCGGTATATTCGAGCCGAAAAAGTCAGATG |
| Y63F RP | TCGGCTCGAATATACCGGGTGTGTCCAG |
| D69A FP | GAAAAAGTCAGCTGTTCTAGGAC |
| D69A RP | CTAGAACAGCTGACTTTTTCGGCTC |
| D69N FP | GAAAAAGTCAAACGTTCTAGGACATTCCATGGTG |
| D69N RP | GTCCTAGAACGTTTGACTTTTTCGGCTCGTA |
| AaEraΔKH FP | TAATAACCCAGCTTTCTTGTACAAAG |
| AaEraΔKH RP | GTACAAGAAAGCTGGGTTATTATCCCTCAGGCAGGTACTTC |

**Table S4: Primers for mutagenesis of *Ec*Era expression vector.**

| Primer | Sequence (5' to 3') |
|---|---|
| H67A FP | GGCCTGGCGATGGAAGAAAAACGCGCCAT |
| H67A RP | TTCCATCGCCAGGCCCGGTGTATCGACGT |
| H67W FP | GGCCTGTGGATGGAAGAAAAACGCGCCAT |
| H67W RP | TTCCATCCACAGGCCCGGTGTATCGACGT |
| H67F FP | GGGCCTGTTCATGGAAGAAAAACGCGCCAT |
| H67F RP | CTTCCATGAACAGGCCCGGTGTATCGACGT |
| H67D FP | GGGCCTGGATATGGAAGAAAAACGCGCCAT |
| H67D RP | CTTCCATATCCAGGCCCGGTGTATCGACGT |
| E69A FP | GCATATGGCGGAAAAACGCGCCATTAACCGCCTG |
| E69A RP | CGTTTTTCCGCCATATGCAGGCCCGGTGTATC |
| E70A FP | ATGGAAGCGAAACGCGCCATTAACCGCCTGATG |
| E70A RP | CGCGTTTCGCTTCCATATGCAGGCCCGGTG |
| E69A.E70A FP | CATATGGCGGCGAAACGCGCCATTAACCGCCTGATG |
| E69A.E70A RP | CGTTTCGCCGCCATATGCAGGCCCGGTGTATCG |
| H67A.E69A.E70A FP | CTGGCGATGGCGGCGAAACGCGCCATTAACCGCCTG |
| H67A.E69A.E70A RP | CGCCGCCATCGCCAGGCCCGGTGTATCGACGTAG |
| M78A FP | CCGCCTGGCGAACAAAGCGGCGAGCAGCTC |
| M78A RP | TTGTTCGCCAGGCGGTTAATGGCGCGTTTTTC |
| M78W FP | CCGCCTGTGGAACAAAGCGGCGAGCAGCTC |
| M78W RP | TTGTTCCACAGGCGGTTAATGGCGCGTTTTTC |
| M78Y FP | CCGCCTGTATAACAAAGCGGCGAGCAGCTC |
| M78Y RP | GCTTTGTTATACAGGCGGTTAATGGCGCGTTTTTC |
| K21A FP | AACGTTGGCGCTTCCACATTGTTGAAC |
| K21A RP | AATGTGGAAGCGCCAACGTTCGGAC |

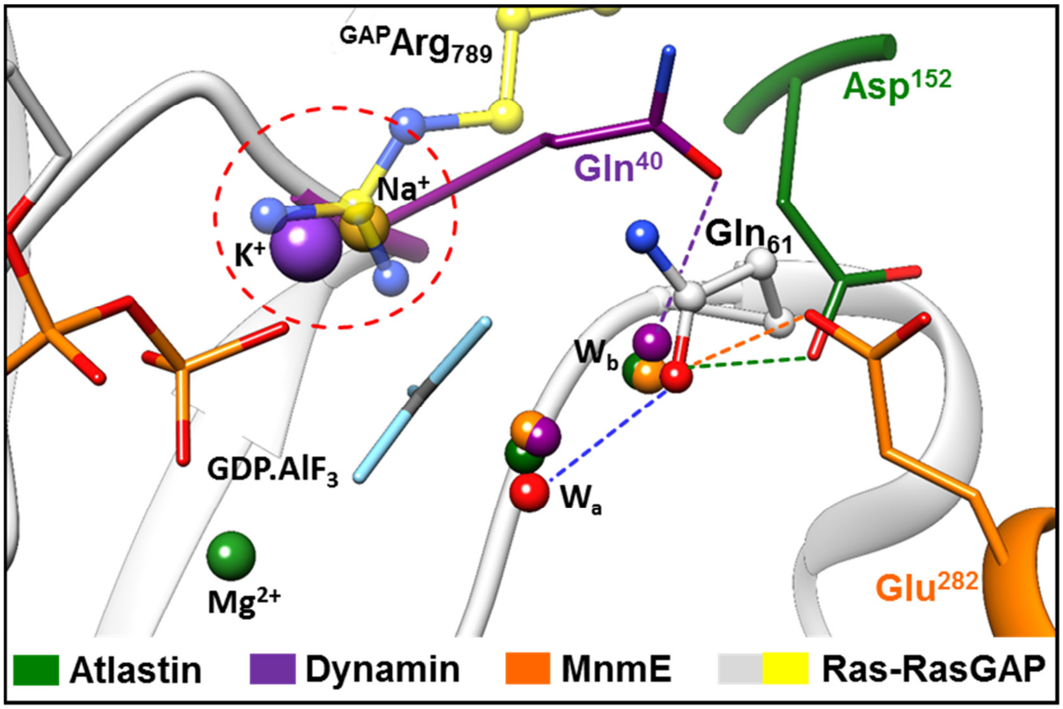


**Figure S1: GTP hydrolysis in HAS-GTPases**

Comparison of active sites of HAS-GTPases: Atlastin (PDB: 4IDO), Dynamin (PDB: 2X2F) and MnmE (PDB: 2GJ8), captured in the presence of transition state analogue GDP.$AlF_x$. The structures are superposed with that of Ras-RasGAP complex (PDB: 1WQ1). Ras is coloured white and the Arginine residue from GAP is coloured yellow. Residues important for catalysis in HAS-GTPases are coloured differently: dark green (Atlastin), purple (Dynamin) and orange (MnmE). Corresponding active site waters are also coloured accordingly, while the attacking water ($W_a$) in Ras is coloured red. The hydrogen bond between $W_a$ and the Gln61 of Ras is shown as a blue dashed line, while those between a bridging water molecule ($W_b$) and corresponding residue from that HAS-GTPase are shown as dashes and coloured accordingly. Cations $K^+$ (violet) and $Na^+$ (golden yellow) that bind in the active sites of MnmE and Dynamin are also shown.

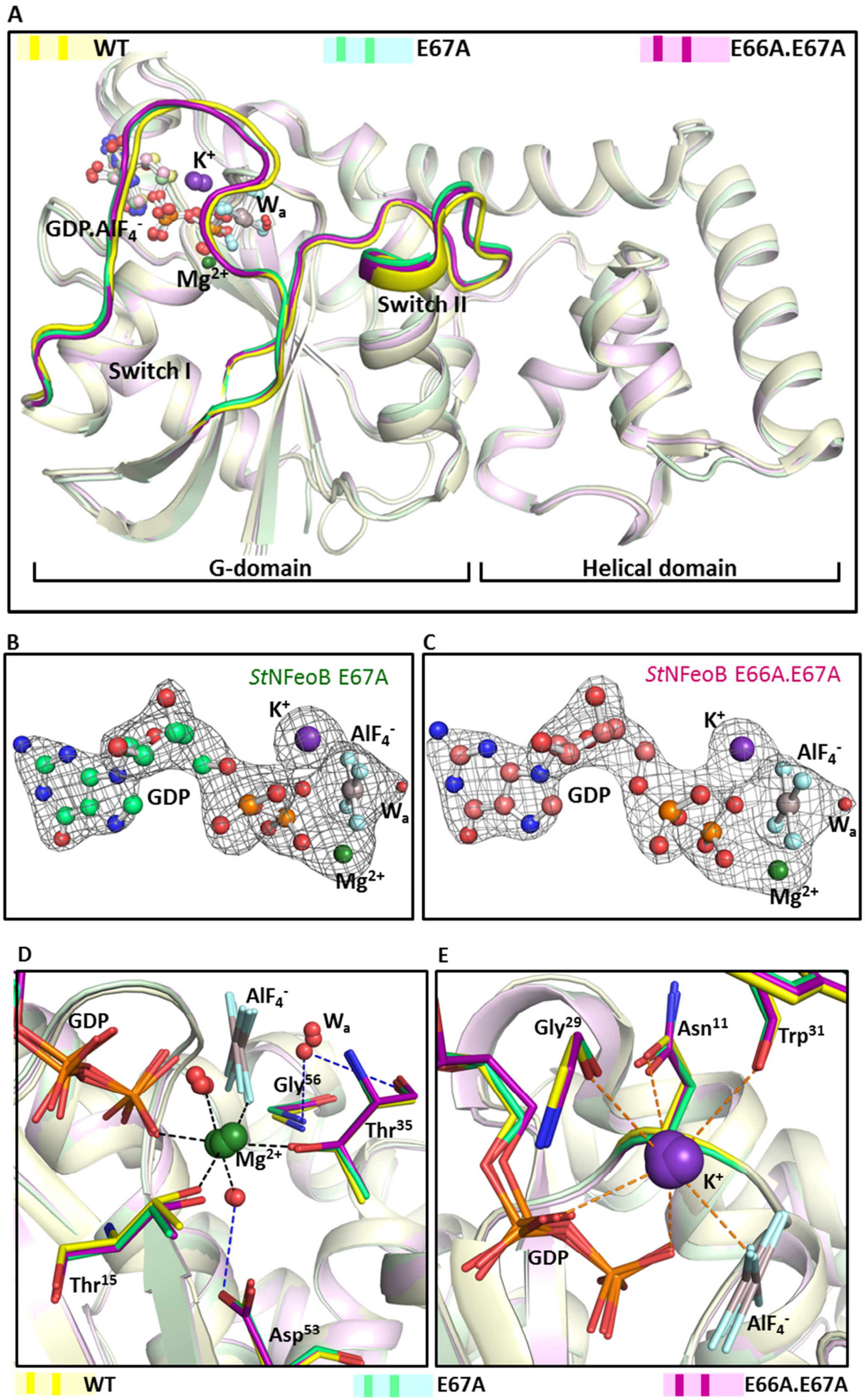
A
WT
E67A
E66A.E67A
K+
Wa
GDP.AlF4-
Mg2+
Switch II
Switch I
G-domain
Helical domain
B
StNFeoB E67A
K+
AlF4-
GDP
Wa
Mg2+
C
StNFeoB E66A.E67A
K+
AlF4-
GDP
Wa
Mg2+
D
AlF4-
Wa
GDP
Gly56
Thr35
Mg2+
Thr15
Asp53
E
Gly29
Asn11
Trp31
K+
GDP
AlF4-
WT
E67A
E66A.E67A

**Figure S2: Overall structures of *St*NFeoB mutants and the active site architecture**

(**A**) Structural superposition of *St*NFeoB WT (yellow) (PDB: 3SS8), E67A (green) and E66A.E67A (purple); switch regions are highlighted in darker shades, respectively. (**B**) & (**C**) The nucleotide-binding site of mutants. The grey mesh represents the Polder omit maps (contoured at 5σ) about the ligands of nucleotide-binding site- GDP.$AlF_4^-$ (transition state analogue), $Mg^{2+}$ (dark green), $K^+$ (violet) and the attacking water ($W_a$) for (**B**) *St*NFeoB E67A (light green) and (**C**) *St*NFeoB E66A.E67A (light pink). The conservation of active site interactions among all the structures (colour scheme as above) that stabilize (**D**) $Mg^{2+}$, $W_{a,}$ and (**E**) $K^+$. The interactions within the WT structure alone have been represented for clarity. The coordinate bonds that stabilize $Mg^{2+}$ (1.8 Å-2.5 Å) are shown as black dashes; the electrostatic interactions with $K^+$ (2.5 Å-3.0 Å) as orange dashes and the hydrogen bonds (3.0 Å-3.5 Å) are represented through blue dashes.

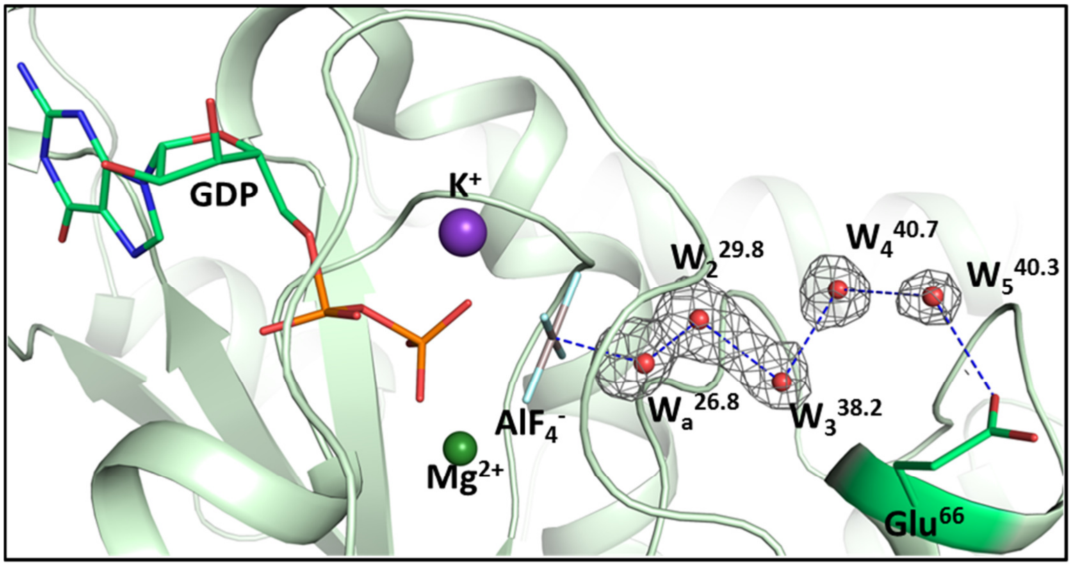


**Figure S3: Active site arrangement of *St*NFeoB E67A**

The transition state structure of *St*NFeoB E67A mutant (light green) illustrating a different arrangement of active site waters and its interaction with Glu66 (green). The waters are labeled as $W_a$ (attacking water) and $W_2$-$W_5$; the value in superscript denotes their respective B-factors. For comparison, the average B-value for the main chain atoms is 38.4. The grey mesh around the water molecules represents the Polder omit map contoured at 5σ. Hydrogen bonds are represented as blue dashes.

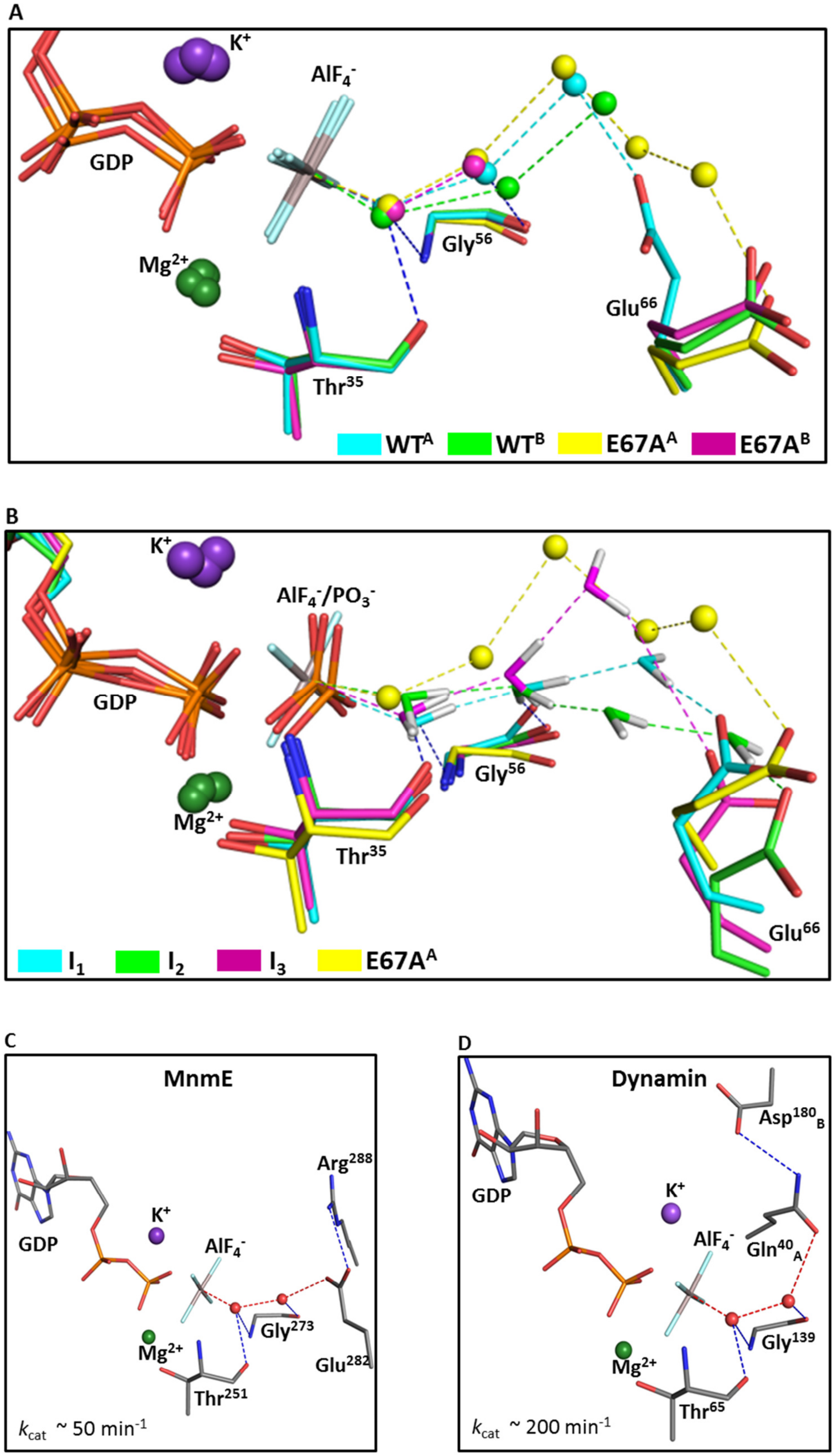
A
K+
AlF4-
GDP
Gly56
Mg2+
Glu66
Thr35
WTA
WTB
E67AA
E67AB
B
K+
AlF4-/PO3-
GDP
Gly56
Mg2+
Thr35
Glu66
I1
I2
I3
E67AA
C
MnmE
Arg288
GDP
K+
AlF4-
Gly273
Mg2+
Glu282
Thr251
kcat ~ 50 min-1
D
Dynamin
Asp180B
GDP
K+
AlF4-
Gln40A
Gly139
Mg2+
Thr65
kcat ~ 200 min-1

**Figure S4: Analysis of active site arrangement of *St*NFeoB E67A**

(**A**) Comparison of active site arrangement among the monomeric chains of *St*NFeoB WT (PDB: 3SS8) and *St*NFeoB E67A (colour scheme shown above). The waters (spheres) and their interactions (dashes) are coloured in accordance with their respective macromolecule chains. Main chain interactions stabilizing the attacking water in the WT structure are shown as blue dashes. (**B**) Comparison of active site arrangement in chain B of *St*NFeoB E67A (yellow) with three different snapshots (colour scheme shown above) from metadynamics simulations of *St*NFeoB ($I_1$-$I_3$), representing the intermediate state of GTP hydrolysis; these were reported previously[24]. The waters are shown as spheres for the crystal structure and as sticks (with hydrogens) for the simulation snapshots. The waters and their interactions (dashes) are represented in the colours of their respective chains. The active site arrangement from transition state structures of HAS-GTPases: (**C**) MnmE (PDB: 2GJ8) and (**D**) Dynamin (PDB: 2X2F). The interactions of GDP.$AlF_4^-$ with the catalytic residue (E282 in MnmE and Q40 in Dynamin) through the chain of water molecules are represented by red dashes; remaining interactions are represented by blue dashes. The $k_{cat}$ values are taken from references 20 and 23. Dynamin exists as a homodimer in the transition state structure, and the residues from different monomers are distinguished through subscripts A and B.

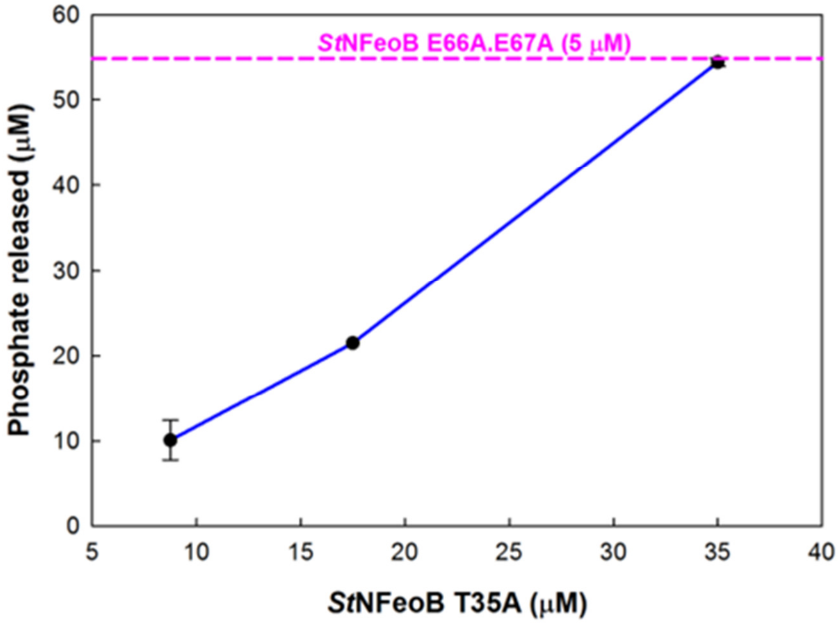


**Figure S5: Catalytic efficiency of *St*NFeoB E66A.E67A**

The extent of GTP hydrolysis after one hour of incubation with different concentrations of T35A (–●–) in comparison to that with 5 μM of E66A.E67A (---).

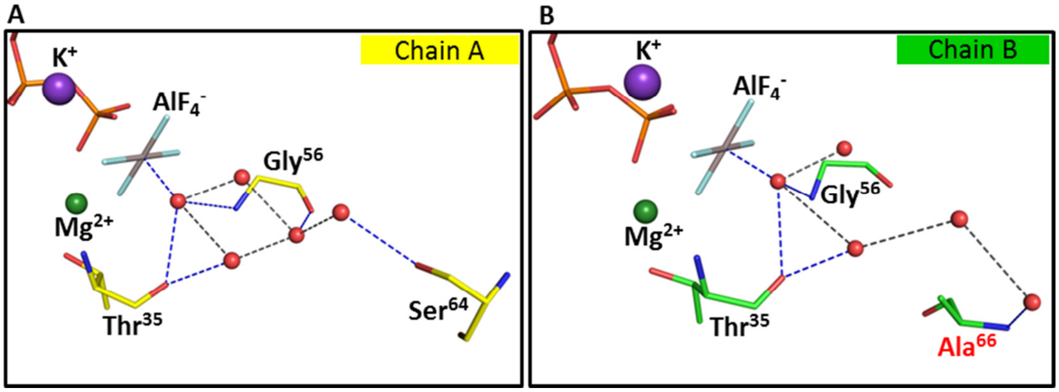


**Figure S6: Arrangement of active site waters in *St*NFeoB E66A.E67A**
The arrangement of active site in (**A**) monomer A (yellow) and (**B**) monomer B (green) of the transition state structure of *St*NFeoB E66A.E67A. Intra-water hydrogen bonds are represented by grey dashes and those with protein by blue dashes. The red-coloured label in (A) highlights the mutated residue.

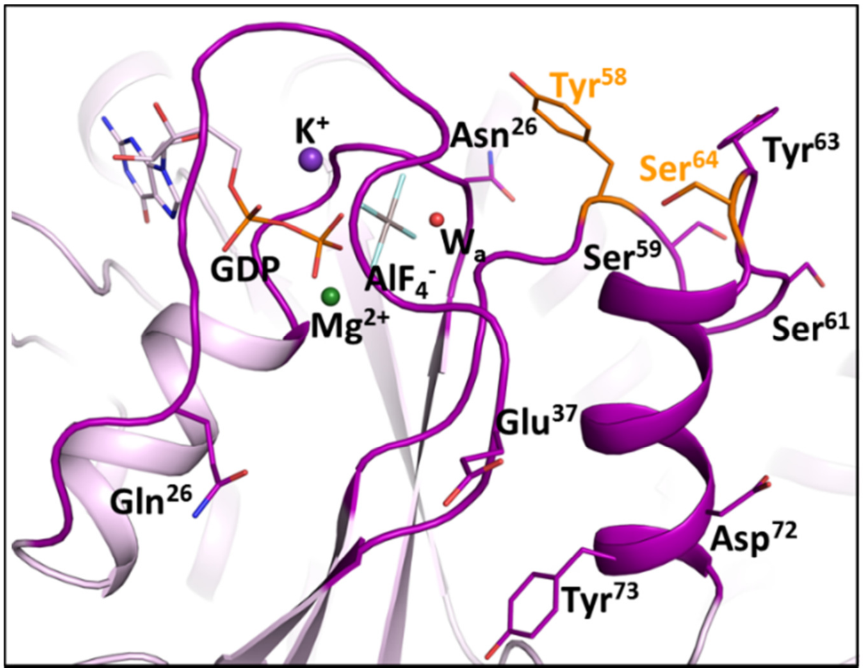


**Figure S7: Analysis of active site of *St*NFeoB E66A.E67A**
The regions located within 15 Å distance of the attacking water ($W_a$) are shown in purple colour. Side chains of polar residues (Asp, Asn, Glu, Gln, His, Ser, Thr and Tyr) in these regions are shown. The residues oriented towards the attacking water are highlighted in orange colour.

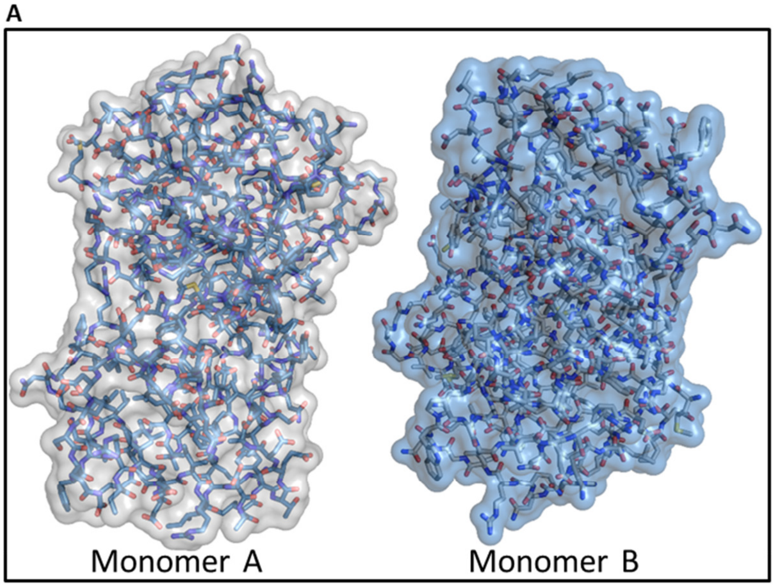


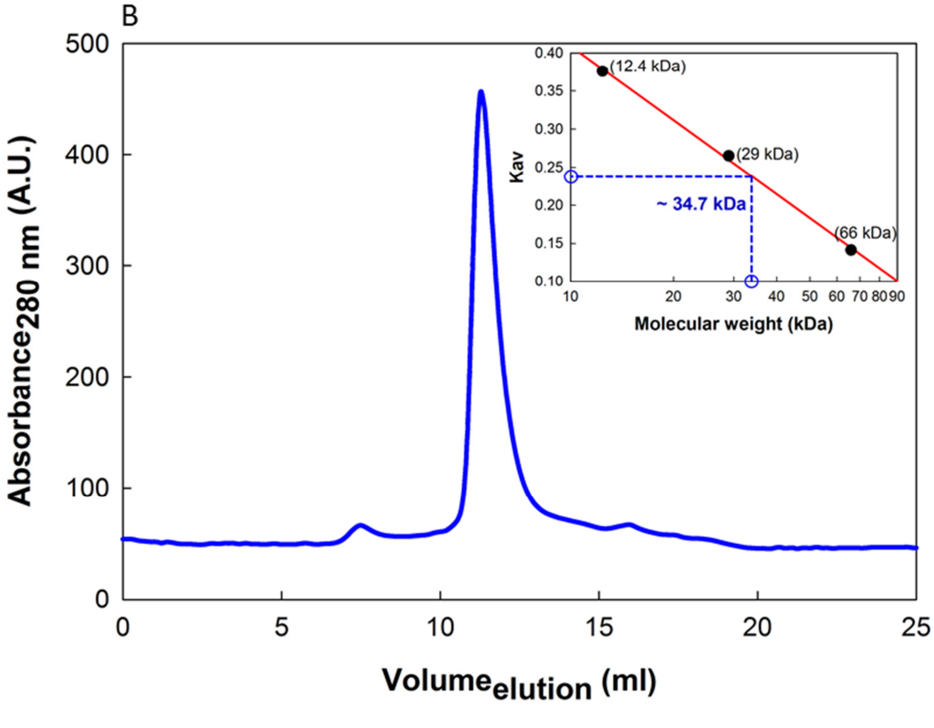


**Figure S8: Analysis of asymmetric unit dimer of *St*NFeoB E66A.E67A**

(**A**) Structure model of the contents of an asymmetric unit of *St*NFeoB E66A.E67A crystal. (**B**) Size exclusion chromatogram of E66A.E67A injected on Superdex75 column, with its calibration curve shown in the inset.

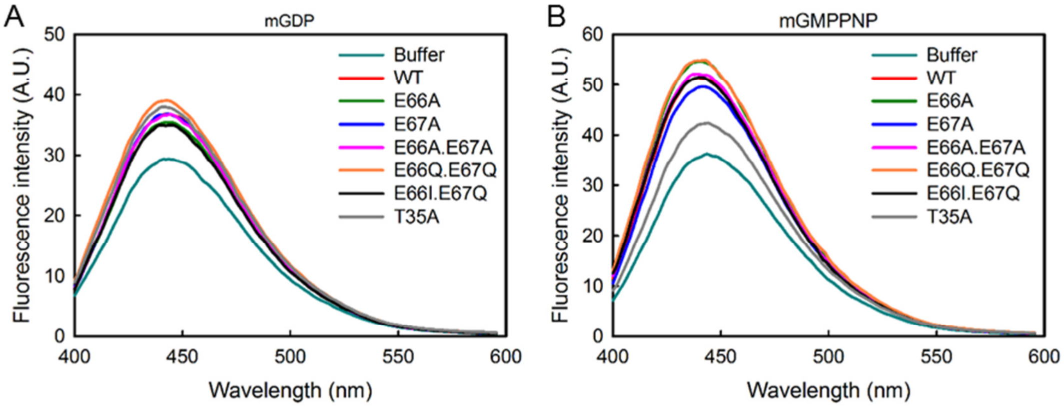


**Figure S9: Nucleotide binding among the *St*NFeoB constructs**

The binding spectra of fluorescent nucleotide: (**A**) mant-GDP and (**B**) mant-GMPPNP (non-hydrolyzable GTP analog) with different proteins (colour scheme indicated above). Here, T35A (grey colour) is taken as a control; this mutation affects the binding of $Mg^{2+}$ and eventually affects the stability of switch-I and binding of GTP.

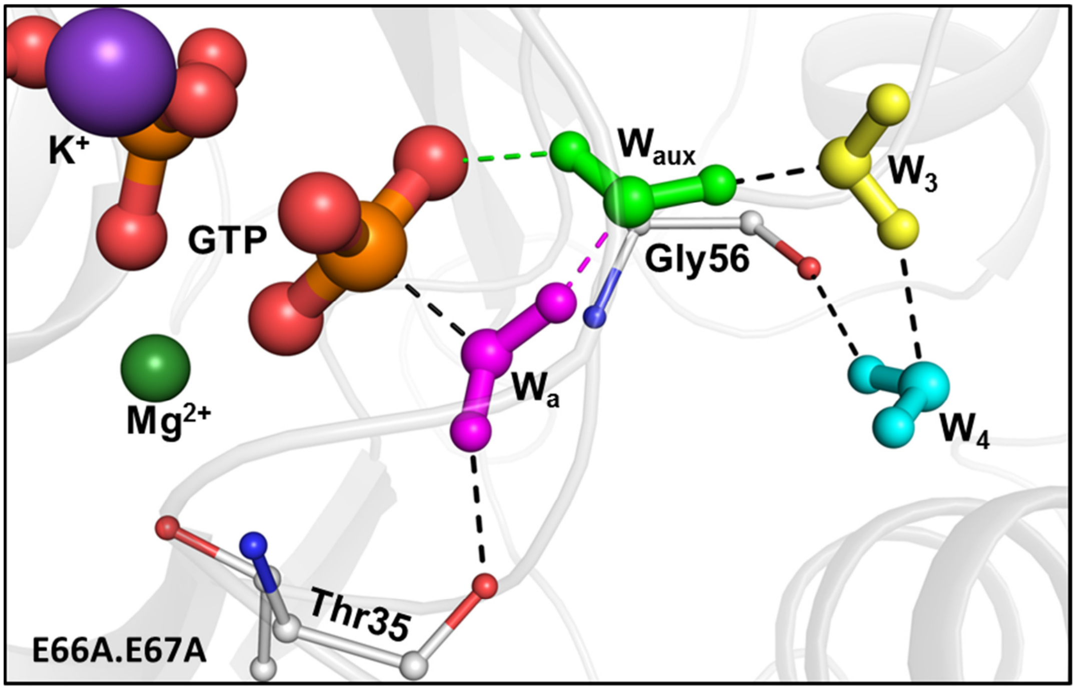


**Figure S10: Transition state of *St*NFeoB E66A.E67A**

Snapshot from QM/MM simulations[24], representing the transition state of GTP hydrolysis in *St*NFeoB E66A.E67A. The waters are coloured differently and represented as ball and sticks with hydrogens. These are labelled $W_a$ (attacking water), $W_{aux}$ (auxiliary water), $W_3$ and $W_4$. The interactions represented as dashed lines, in colours of respective water molecules, indicate the direction of proton transfer; other interactions are represented as black dashes.

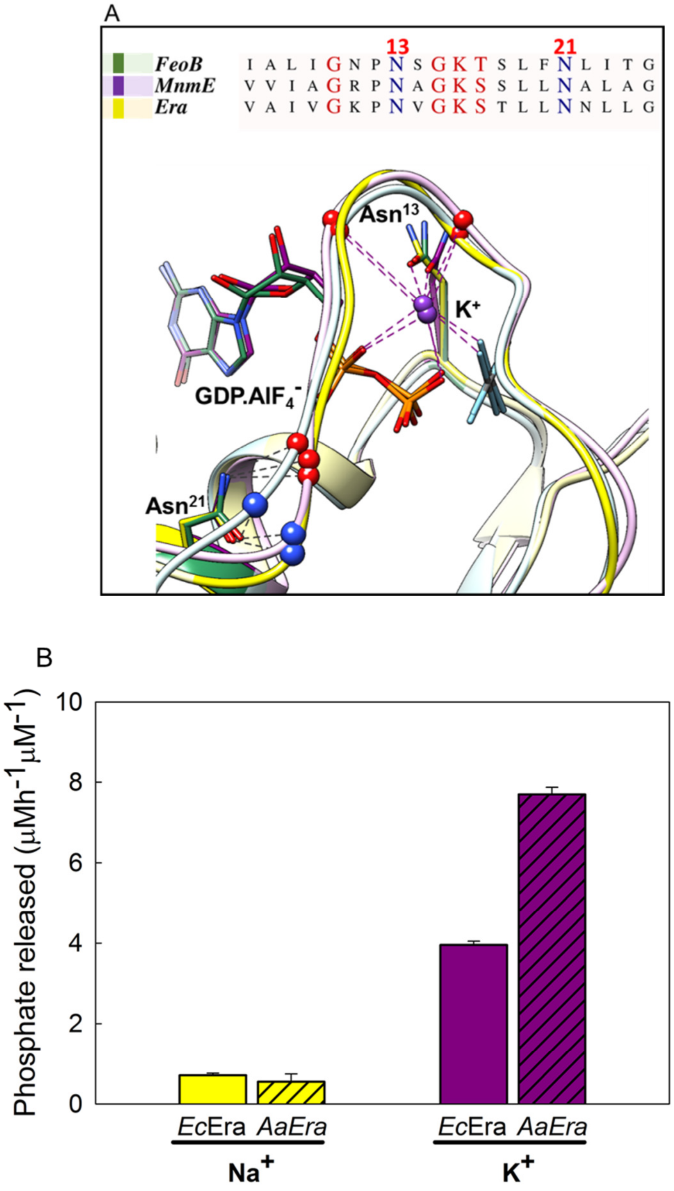


**Figure S11: Stimulation of GTP hydrolysis of Era**

**(A)** Sequence and structural comparison of $K^+$ binding regions of *Aa*Era (PDB: 3IEV) with $K^+$ dependent GTPases: FeoB (PDB: 3SS8) and MnmE (PDB: 2GJ8). Electrostatic interactions with $K^+$ are represented as purple dashes and hydrogen bonds as grey dashes; the main chain atoms are shown as spheres on the cartoon ribbon. **(B)** Comparing $Na^+$ and $K^+$ in stimulating the GTP hydrolysis of Era.

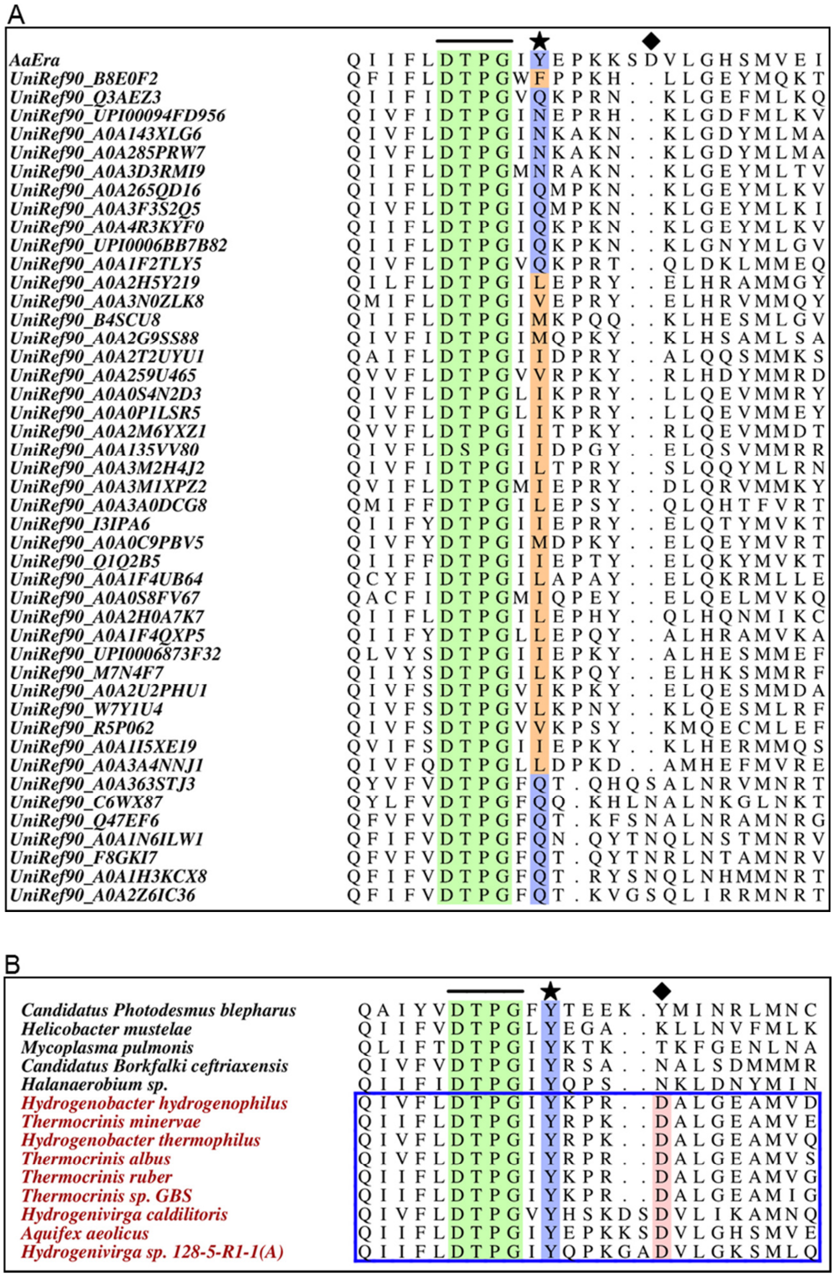


**Figure S12: Sequence alignments of Era homologues**

**(A)** Sequences from *Consurf* analysis of Era homologues that show variations in the position corresponding to Y63 of *Aa*Era in spite of strong conservation score for the position. **(B)** Alignment of all the Era homologues that have Y at the position corresponding to residue 63 of *Aa*Era. Sequences enclosed in the blue box belong to *Aquificae* phylum. In both the alignments, the G3 motif is marked by a line, position corresponding to residue 63 by a star symbol, and position corresponding to residue 69 is marked by a diamond symbol.

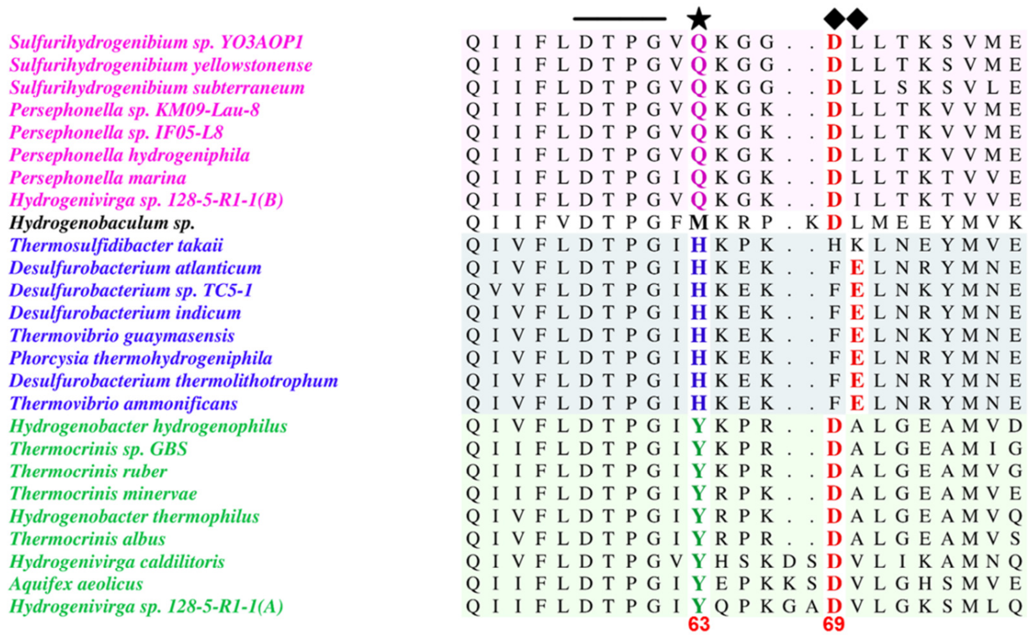


**Figure S13: Sequence alignments of Era homologues from *Aquificae* phylum**

Alignment of all the Era homologues belonging to *Aquificae* phylum. G3 motif is marked by a line, position corresponding to residue 63 by a star symbol, and the strong conservation of an acidic residue at the position corresponding to residue 69 or at adjacent position is marked by a diamond symbol.

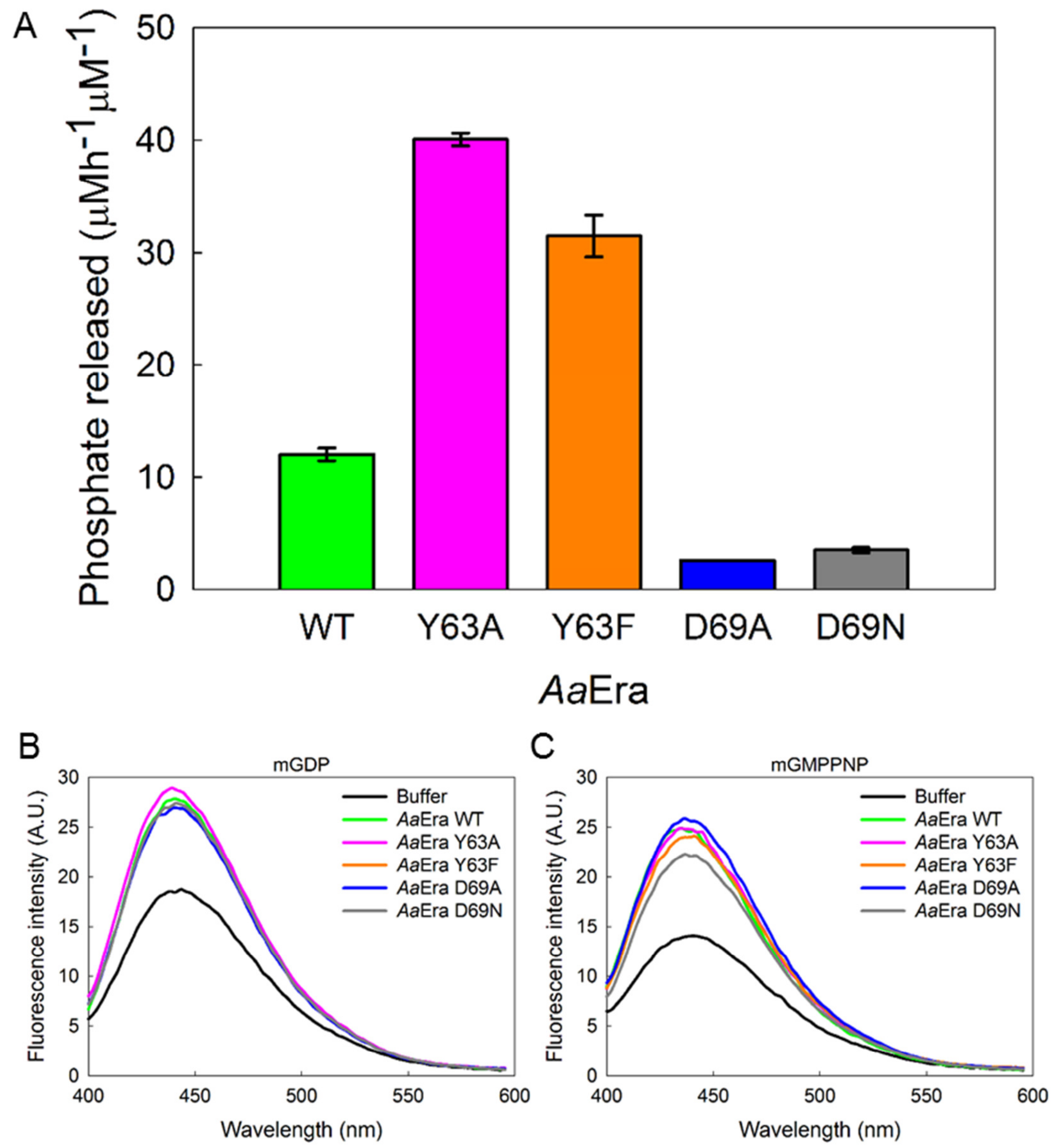


**Figure S14: GTP hydrolysis of *Aa*Era constructs**

**(A)** Comparison of GTP hydrolytic activities of different *Aa*Era mutants. **(B)** and **(C)** Fluorescently labeled-nucleotide binding spectra of *Aa*Era constructs with mant-GDP or mant-GMPPNP, respectively.

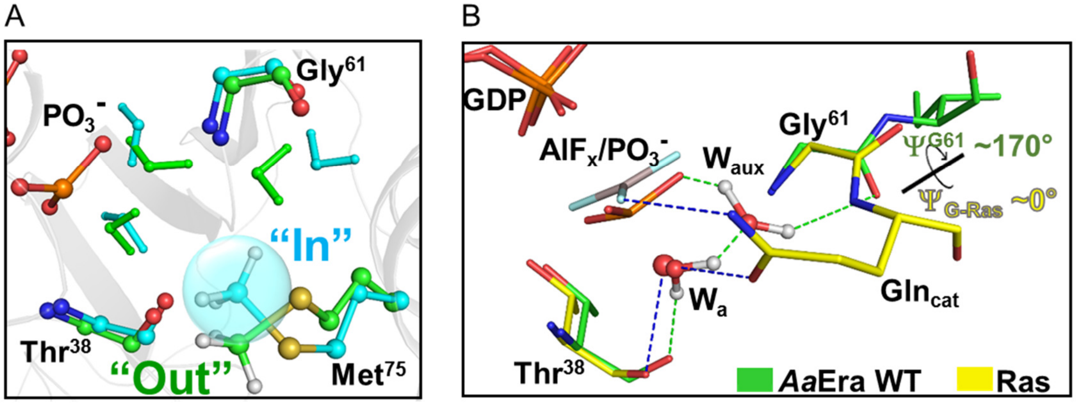


**Figure S15: GTP hydrolysis in *Aa*Era**

**(A)** Comparison of active site arrangement of *Aa*Era WT in GTP bound state (cyan) *v/s* transition state of GTP hydrolysis[25] (green). **(B)** Comparison of active site arrangement in transition state snapshot of *Aa*Era WT (green) with the structure of Ras (yellow) bound to transition state analogue (PDB: 1WQ1). The attacking water is shown as red spheres for the crystal structure and as ball and sticks for the simulation snapshot. The hydrogen bonds are shown as dashes of green and blue colours for *Aa*Era WT and Ras, respectively.

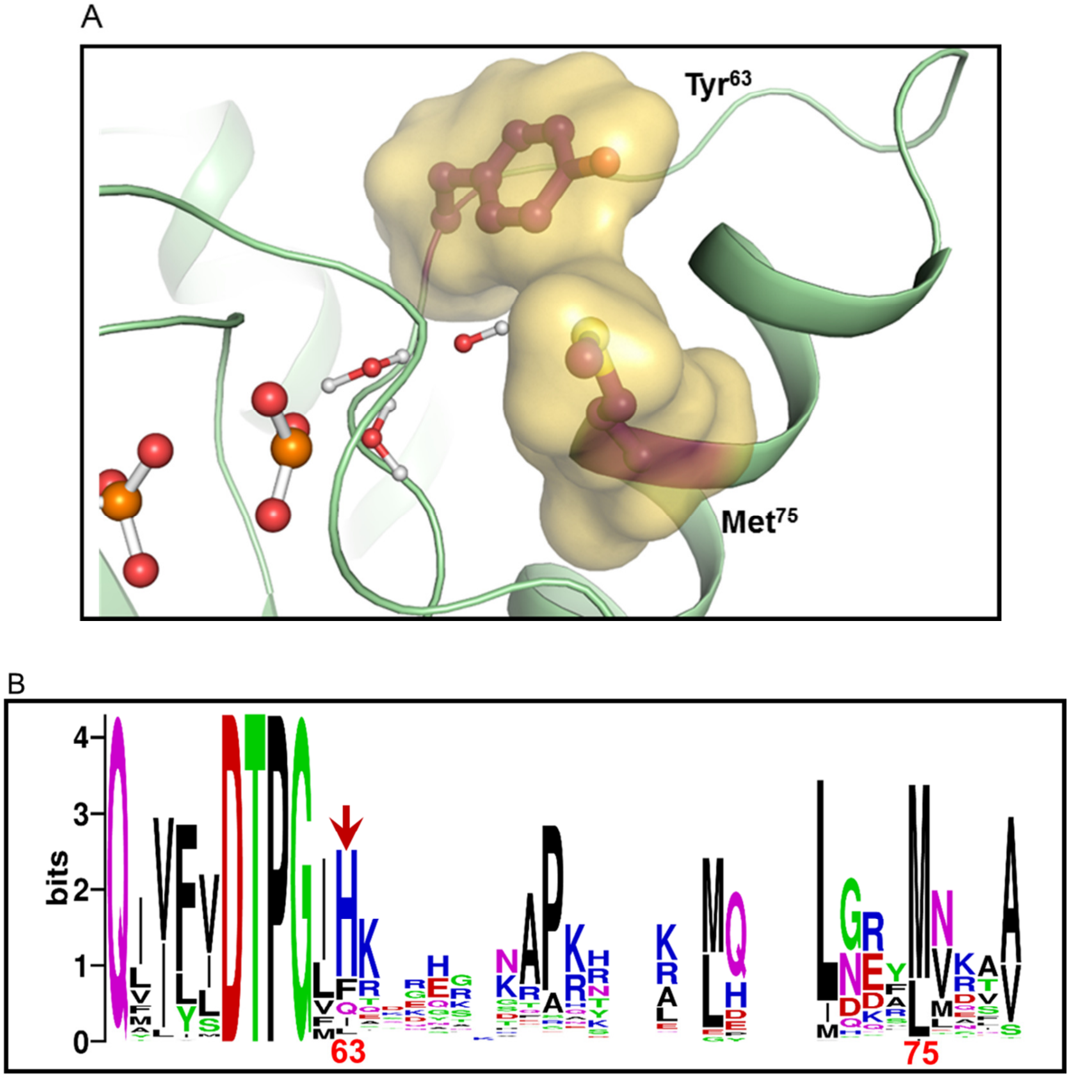


**Figure S16: Structure and sequence analysis of Era**

**(A)** Surface representing the interface between Y63 and M75 in the transition state snapshot of GTP hydrolysis in *Aa*Era WT QM/MM simulation[25]. **(B)** Sequence logo representation (generated using *WebLogo*) of the aligned sequences of Era homologues.

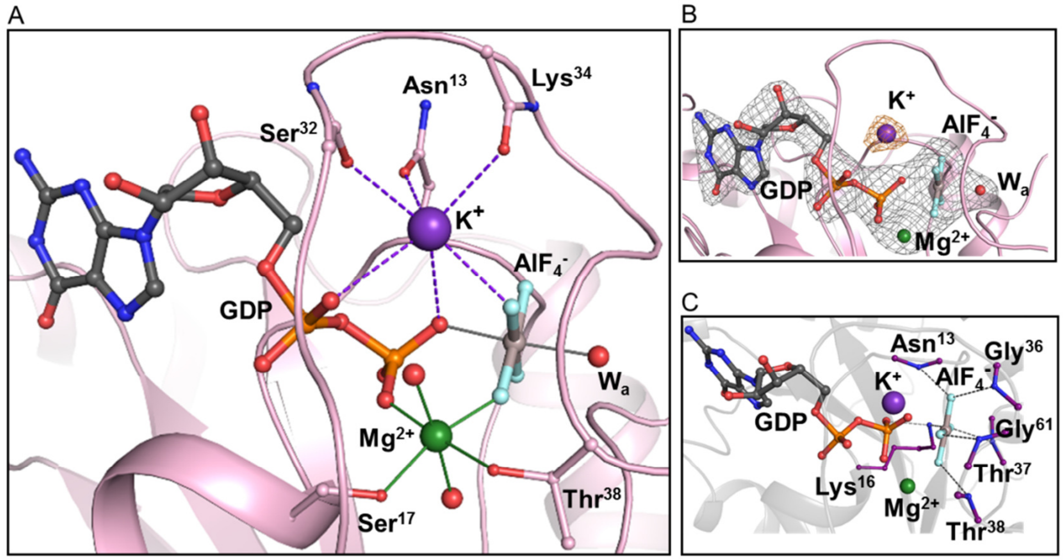


**Figure S17: Active site arrangement of *Aa*Era Y63A**

**(A)** Interactions of active site ions in the *Aa*Era Y63A structure. The electrostatic interactions (2.5 Å-3.0 Å) that stabilize $K^+$ are shown with violet-coloured dashed lines. The coordinate bonds (1.8 Å-2.5 Å) of $Mg^{2+}$ and $Al^{3+}$ (of $AlF_4^-$) are represented through solid lines of green and grey colours, respectively. **(B)** Polder omit map around the ligands – GDP.$AlF_4^-$ transition state analogue, magnesium ion ($Mg^{2+}$) and attacking water ($W_a$), and anomalous-difference LLG Fourier map around potassium ion ($K^+$) bound in the nucleotide-binding site of *Aa*Era Y63A. The grey mesh represents the Polder omit map (contoured at 5σ) and orange mesh depicts the likelihood-based anomalous difference map indicating the position of $K^+$ (contoured at 3.5σ). **(C)** Active site interactions provided by side-chain of K16 and main-chain amides that stabilize the negatively charged $AlF_4^-$.

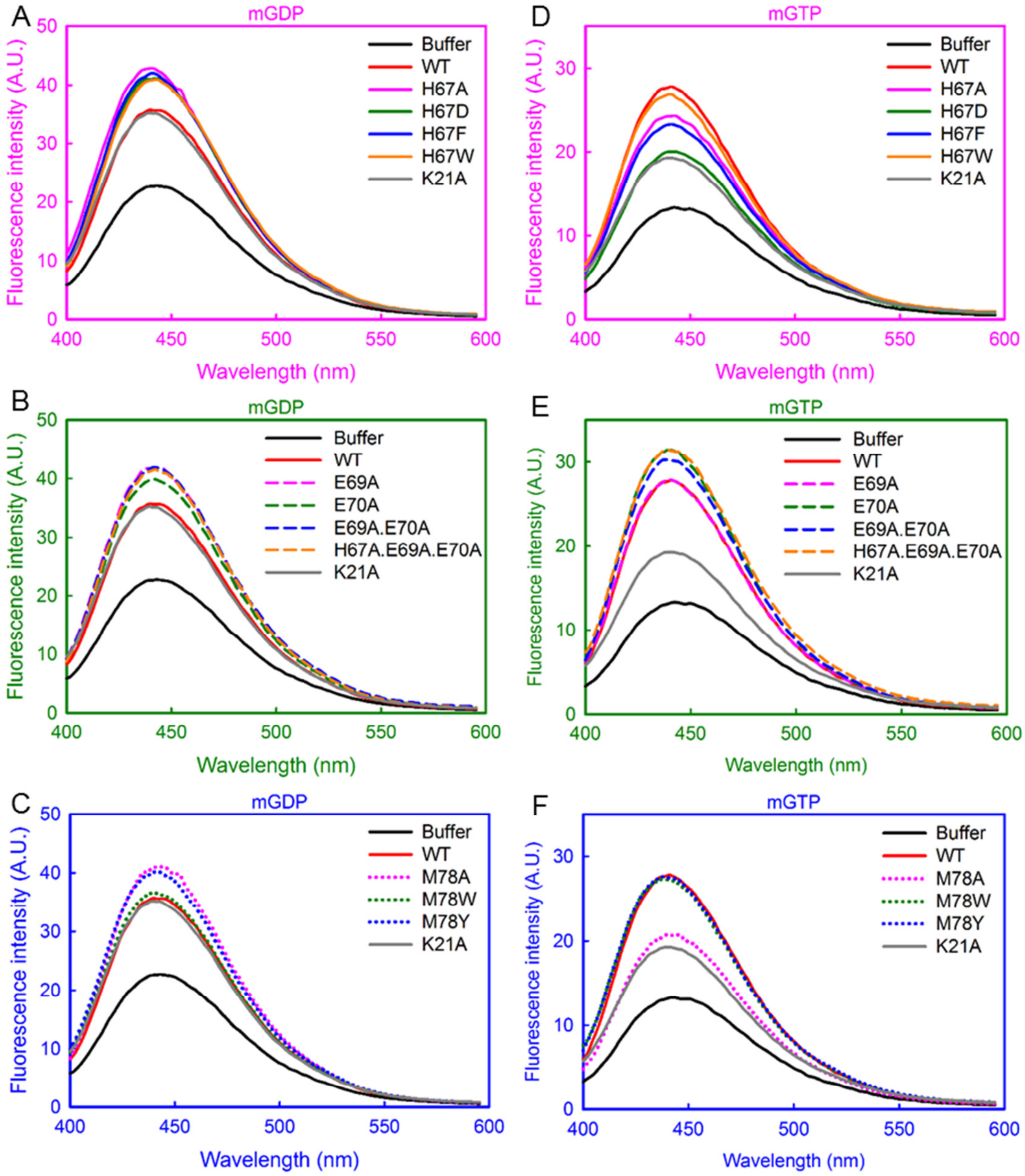


**Figure S18: Nucleotide binding among *Ec*Era constructs**

Fluorescently labelled-nucleotide binding spectra of *Ec*Era constructs with mant-GDP [(**A**)-(**C**)] or mant-GTP [(**D**)-(**F**)], respectively. The frames of plots are coloured differently to segregate constructs with substitutions at a particular locus for convenient representation of data.
